# Selecting for altered substrate specificity reveals the evolutionary flexibility of ATP-binding cassette transporters

**DOI:** 10.1101/797100

**Authors:** Sriram Srikant, Rachelle Gaudet, Andrew W. Murray

**Affiliations:** Department of Molecular and Cellular Biology, Harvard University

## Abstract

ABC transporters are the largest family of ATP-hydrolyzing transporters, with members in every sequenced genome, which transport substrates across membranes. Structural studies and biochemistry highlight the contrast between the global structural similarity of homologous transporters and the enormous diversity of their substrates. How do ABC transporters evolve to carry such diverse molecules and what variations in their amino acid sequence alter their substrate selectivity? We mutagenized the transmembrane domains of a conserved fungal ABC transporter that exports a mating pheromone and selected for mutants that export a non-cognate pheromone. Mutations that alter export selectivity cover a region that is larger than expected for a localized substrate-binding site. Individual selected clones have multiple mutations which have broadly additive contributions to specific transport activity. Our results suggest that multiple positions influence substrate selectivity, leading to alternative evolutionary paths towards selectivity for particular substrates, and explaining the number and diversity of ABC transporters.

## Introduction

During evolution many genes have duplicated and diverged to acquire new functions. ABC transporters are an extreme example of protein diversification because they represent the largest family of ATP-hydrolyzing transporters that has descended from a common origin; every sequenced cellular genome contains at least one member of this family. ABC transporters use ATP hydrolysis to drive substrate import or export (Ford and Beis, 2019). Most genomes contain paralogous transporters with different family members transporting different substrates and playing different physiological roles. ABC transporters contain cytoplasmic nucleotide binding domains (NBD) with conserved motifs for binding and hydrolyzing ATP, connected to transmembrane domains (TMD) that undergo conformational changes to transport substrates across membranes (Srikant and Gaudet, 2019).

ABC exporters transport substrates from the cytosol to the extracellular or luminal compartment. All eukaryotic exporters belong to a homologous subfamily called the type I ABC exporters and perform crucial physiological processes by exporting substrates with a wide variety of physicochemical properties. For example, type 1 exporters include MsbA, which transports a large glycosylated lipid (lipopolysaccharide (LPS)) for bacterial outer membrane biogenesis (Mi et al., 2017; Tefsen et al., 2005), Atm1, which exports a mitochondrial compound needed to form iron-sulfur (Fe-S) clusters in eukaryotic cytosolic proteins (Kispal et al., 1999; Srinivasan et al., 2014), and the Transporter for Antigen Processing (TAP), which exports the peptides that are presented to the adaptive immune system of vertebrates from the cytosol to the endoplasmic reticulum (Abele and Tampe, 2018). The substrate selectivity of ABC exporters determines the multidrug resistance phenotype of cancers (e.g. P-glycoprotein or P-gp;(Robey et al., 2018)), and parasitic pathogens (*P. falciparum* MDR1; (Koenderink et al., 2010)). Sequence conservation in the TMDs of homologous exporters has not been helpful in identifying a conserved binding site for cognate substrates in orthologous or paralogous exporters. Structures of inward-open exporters with different substrates are remarkably similar, with substrate expected to bind over the large, chemically heterogenous surface of the TMD cavity (Srikant and Gaudet, 2019). Biochemical crosslinking and mutagenesis experiments on P-gp and TAP have identified positions in the TMD cavity involved in substrate-recognition (Geng et al., 2015; Loo et al., 2003; Loo and Clarke, 2000), but a system to characterize the sequence determinants of substrate selectivity in type I ABC exporters has not been reported.

To study substrate selectivity in ABC exporters we took advantage of the pheromones used for fungal mating. The Dikarya is the most species-rich and familiar group of fungi, spanning about a billion years of evolution, and is split into two groups, the ascomycetes, which includes the baker’s yeast, *Saccharomyces cerevisiae*, and the basidiomycetes. In Dikarya, mating depends on the exchange of diffusible peptide pheromones between the two haploid mating partners (gametes; Figure 1a). The first identified fungal pheromone, RhodotorucineA (Kamiya et al., 1978), is a farnesylated peptide produced by a basidiomycete yeast, *Rhodosporidium toruloides.* Basidiomycetes use only farnesylated pheromones whereas Ascomycetes like *S. cerevisiae* use a farnesylated peptide in one mating type and an unmodified peptide pheromone in the other. Genetic analysis in *S. cerevisiae* identified the farnesylated pheromone, **a**-factor (Anderegg et al., 1988; Michaelis and Herskowitz, 1988), which is synthesized in the cytosol of *MAT***a** haploid cells (**a**-cells) and exported into the medium by a type I ABC exporter, Ste6 (Mackay and Manney, 1974). The restriction of Ste6’s function and expression to **a**-cells (Kuchler et al., 1989; Wilson and Herskowitz, 1986) and biochemical assays of **a**-factor transport (Ketchum et al., 2001; Michaelis, 1993) strongly argue for Ste6 being a dedicated **a**-factor (farnesylated pheromone) exporter. With the known variation in the peptide sequence of farnesylated pheromones across fungal phylogeny (Caldwell et al., 1995), we expect coevolution of substrate selectivity in the homologous pheromone exporters. Because the physiologically relevant substrates evolve across the Dikarya, we hypothesized that the Ste6 family of pheromone exporters would coevolve with the pheromones, allowing us to study the variation of substrate selectivity.

**Figure 1.**
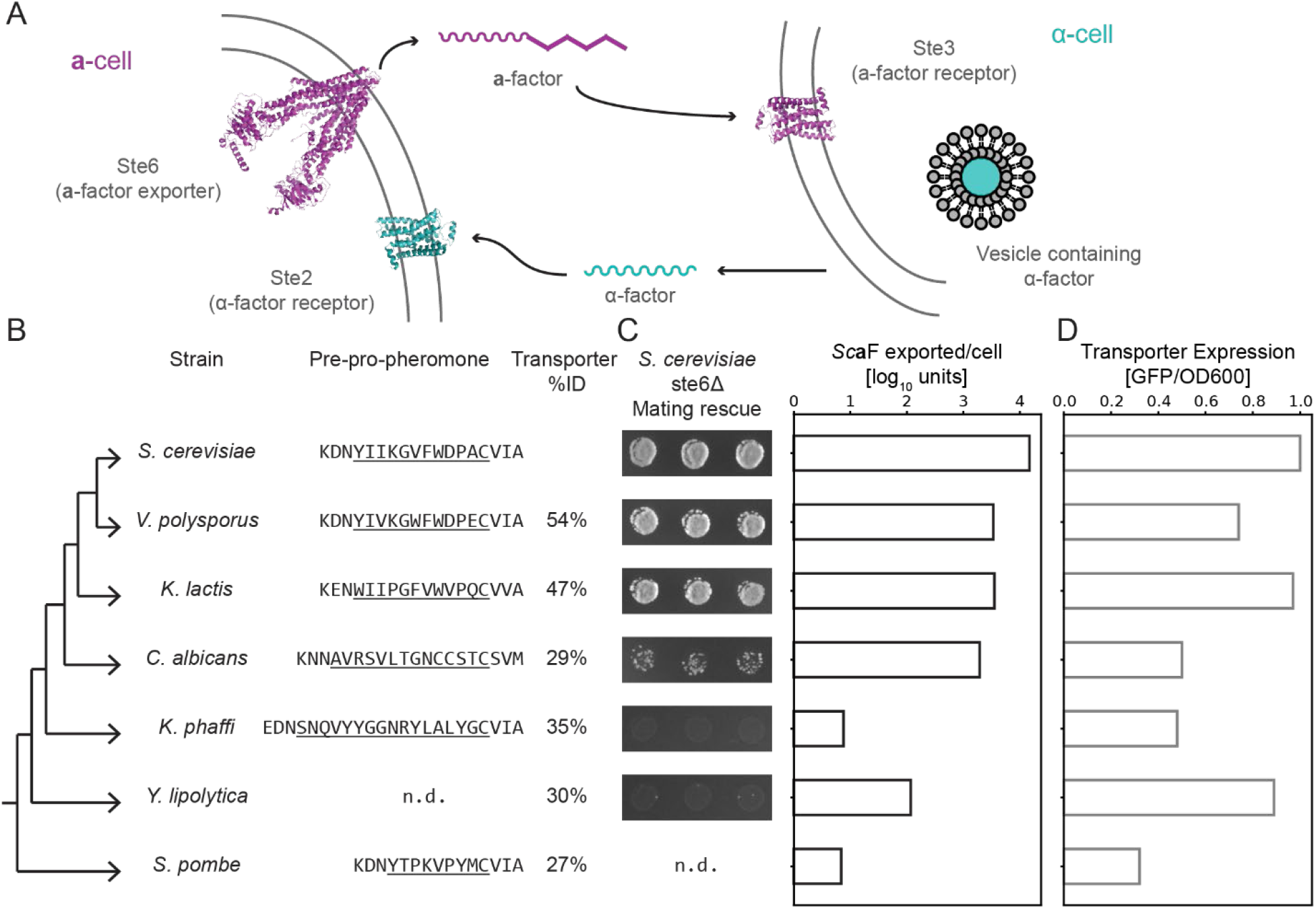
Homologous pheromone transporters can partially rescue a-factor export from *S. cerevisiae*. (**A**) Mating in fungi is initiated by a two-way pheromone communication between haploids of different mating types. In *S. cerevisiae*, α-cells secrete the 13-amino acid peptide α-factor (αF) via secretory vesicles; αF is recognized by a G-protein coupled receptor (GPCR), Ste2, on **a**-cells. The farnesylated 12-amino acid, **a**-factor, is made in the cytoplasm of **a**-cells exported by a dedicated ABCB exporter, Ste6, and recognized by a different GPCR, Ste3, on α-cells. (**B**) A cladogram of the yeasts used in this study, highlighting the sequence of **a**-factor-like pheromones (mature peptides are underlined and only a portion of the N terminal region of the initial peptide is shown; (Davey, 1992; Dignard et al., 2007; Heistinger et al., 2018; Michaelis and Herskowitz, 1988)) and the % identity of orthologous pheromone transporters to *S. cerevisiae* Ste6. (n.d. is “not determined”.) (**C**) Bioassays to measure **a**-factor export from *ste6Δ S. cerevisiae* **a**-cells that express orthologous pheromone transporters (the transporter sources are the species aligned in (b)). Left: **a**-factor export measured indirectly via mating rescue. Right: **a**-factor export measured by purifying **a**-factor from culture and quantifying its ability to arrest α-cells (**D**) The expression of Ste6 homologs, measured by the per-cell fluorescent signal of cells expressing GFP-tagged transporters.

We started by testing the ability of Ste6 homologs from different Ascomycetes to rescue *S. cerevisiae* **a**-factor export. The Ste6 homolog from *Y. lipolytica*, which last shared a common ancestor with *S. cerevisiae* 320 Mya, exports **a-**factor poorly, allowing us to develop a high-throughput assay for mutations in this protein that increased its ability to export **a**-factor from *S. cerevisiae.* Using libraries of mutagenized transporters, we identified regions of the TMD that affect substrate selectivity by allowing transport of this “novel” substrate. We hypothesize that the large target size for mutations that improve **a**-factor export creates many alternative paths for the evolution of paralogous exporters and explains the success of the ABC transporter family in evolving to “solve” the problem of transporting different substrates.

## Results

### The pheromone and transporter form a conserved module in the mating systems of ascomycetes

Pheromones of many yeasts (unicellular fungi) have been identified (Davey, 1992; Dignard et al., 2007; Heistinger et al., 2018; Michaelis and Herskowitz, 1988) with their peptide sequences varying across phylogeny. In one mating type, these pheromones are small peptides that are C-terminally farnesylated and methyl-esterified (Figure 1b); they undergo maturation through a conserved set of enzymes, highlighting the ancestral role of farnesylated pheromones in fungal mating (Chen et al., 1997).

The discovery of orthologs of the Ste6 transporter in *Schizosaccharomyces pombe* (Christensen et al., 1997) and *Candida albicans* (Raymond et al., 1998) validates using homology search algorithms (such as BLAST or HMMER) as a reliable method to identify orthologs of this protein in all sequenced Ascomycete genomes (see Figure 1b). By sequence homology, these transporters are classified as ABCB exporters (Decottignies and Goffeau, 1997). They are most similar to each other in their cytoplasmic ATP-binding domains (NBDs) and show significant variation in the transmembrane domains (TMDs), where substrates are expected to bind. To test the function of homologous pheromone transporter pairs from fungi, we started with a set of lab yeasts whose mating system has been studied and are estimated to have last shared a common ancestor roughly 320 million years ago (Shen et al., 2018).

In *S. cerevisiae*, Ste6 is expressed only in *MAT***a**-cells and *ste6Δ* cells (and those carrying ATP hydrolysis-deficient mutations) are mating deficient (Kuchler et al., 1989), providing a biological assay for **a**-factor transport. We expressed Ste6 orthologs from other species in *ste6Δ S. cerevisiae MAT***a**-cells and used a mating rescue experiment to show that orthologous transporters have functionally diverged (Figure 1c-d). The efficiency of mating roughly correlates with the phylogenetic distance between *S. cerevisiae* and the yeast containing the supplied the Ste6 ortholog, which is consistent with coevolution of the transporter with the pheromone it transports (Figure 1b). To confirm that the efficiency of mating is a good measure of transporter function, we assayed pheromone export directly by collecting exported **a**-factor from growing cultures and using a serial dilution bioassay to measure the quantity of exported pheromone (Figure 1c). We also measured the expression and location of Ste6 orthologs by tagging them with GFP (Figure 1d). A Ste6 ortholog that was strongly expressed but strongly deficient in **a**-factor export is an ideal substrate to mutate and select for increased export of *S. cerevisiae* **a**-factor. Because the *Y. lipolytica* Ste6 (*Yl*Ste6) and *S. cerevisiae* Ste6 (*Sc*Ste6) are equally strongly expressed and show a 100-fold difference in transport of *Sc***a**-factor, we set out to identify mutations in *Yl*Ste6 that would increase the transport of *Sc***a**-factor and shed light on the evolution of substrate specificity in the pheromone transporter family.

### Building a selection system for a-factor transport

We wanted to use a genetic selection to find mutations that increase transport function because it allows us to start with a large, unbiased library of variants. Pheromone export is difficult to select for directly because “success” involves the transport of pheromone from the cytosol to the extracellular medium and **a**-factor is highly hydrophobic, adsorbing to glass and plastic. We thus designed an indirect selection, in which a cell responds to the pheromone that it has exported instead of responding, as cells normally do, to a pheromone from cells of the opposite mating type. This scheme depends on two properties of the pheromone response. First, in *S. cerevisiae* the signaling cascade downstream of the G-protein coupled pheromone receptors—the **α**-factor receptor Ste2 in **a**-cells and the **a**-factor receptor Ste3 in **α-**cells—is the same (Alvaro and Thorner, 2016), meaning that the **a**-factor receptor Ste3 can be expressed and function in an **a**-cell. Thus by replacing Ste2 with Ste3 in a-cells, we engineered an **a**-cell that can detect the pheromone it has exported. Second, pheromone-induced promoters can drive the expression of fluorescent proteins as a reporter for the pheromone-induced signaling cascade (Poritz et al., 2001). To allow us to turn the pheromone response off after each round of selection, we expressed the **a**-factor receptor conditionally from the *S. cerevisiae GAL1* promoter (*P_GAL1_*). Transferring cells from galactose-to glucose-containing medium eliminates receptor expression and allows cells to recover from pheromone-stimulated cell cycle arrest.

Figure 2a shows the design of our selection strain: expressing the **a**-factor receptor, Ste3, in **a**-cells with a pheromone-stimulated reporter (P_FUS1_-ymNeonGreen) creates a cell whose response to the pheromone it exports can be measured via whole-cell fluorescence. By tagging the transporter with an orthogonal fluorescent protein (ymKate2), we created a two-color autocrine system that simultaneously reports on the expression of the transporter and the response to the exported **a**-factor, allowing us to assess the specific activity of the transporter. We tested this system by assaying cells expressing either *Sc*Ste6 or *Yl*Ste6 with flow cytometry to measure the dynamic range of the autocrine reporter (Figure 2b) and confirmed that *Yl*Ste6 was defective in *Sc***a**-factor export. By including bovine serum albumin (BSA), which binds tightly to hydrophobic substances (like the farnesyl group in **a**-factor) in solution, we prevented **a**-factor exported from one cell from stimulating neighboring cells who were unable to export **a**-factor (Figure 2-figure supplement 1). Briefly, cells expressing *Sc*Ste6 and *Yl*Ste6 were mixed at [1:1] or [1:10], respectively, and sorted (Fluorescence-Assisted Cell Sorting, FACS) by gating on autocrine signal that was greater than the brightest 1% of *Yl*Ste6 population. Sorted cells from the mixed populations were identified by unique genetic markers expressed by the *Sc*Ste6 and *Yl*Ste6 strains. Our FACS enrichment protocol leads to a 20-fold enrichment of cells expressing *Sc*Ste6 over those expressing *Yl*Ste6. This test reveals that we can use this system to enrich pooled libraries of mutant transporters (*Yl*Ste6) for clones that export *Sc***a**-factor better by selecting for cells that most strongly express the autocrine reporter.

**Figure 2.**
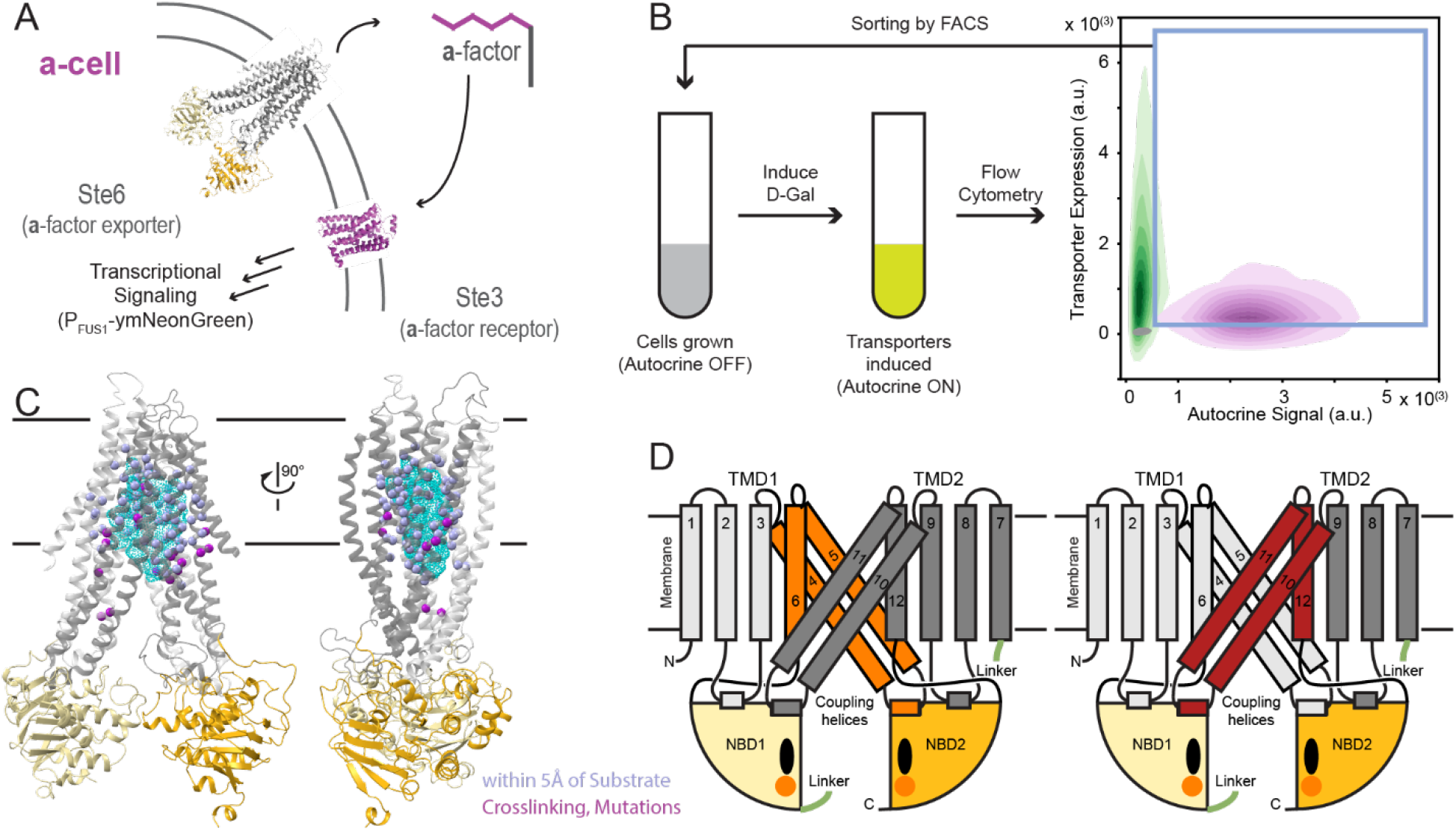
Two-color autocrine system allows for selection of cells expressing transporters functional in the export of *S. cerevisiae* a-factor. (**A**) An autocrine system was designed in **a**-cells, by expressing the **a**-factor sensitive GPCR (Ste3) and a reporter for pheromone stimulation (*P_FUS1_-mNeonGreen*) (Poritz et al., 2001). (**B**) Schematic of a round of enrichment for increased **a**-factor secretion. A period of replicative growth with the autocrine system OFF is followed by FACS with after turning the autocrine reporter ON. Cells were gated on both transporter expression and autocrine signal (blue box) and sorted into autocrine OFF media for expansion before the next round of selection. The 2D histogram shows the flow cytometry signal for populations of cells expressing *Sc*Ste6 (magenta), *Yl*Ste6 (green), and Autocrine OFF *Yl*Ste6 (grey). Populations (n > 25,000 events) are plotted as density contours of autocrine reporter signal and transporter expression signal. (**C**) Model of *Yl*Ste6 (in its inward-open conformation) highlighting positions (Cα as spheres) proximal to substrate density (light blue) in structures of three ABC proteins (P-gp (Zosuquidar in 6FN4 and QZ-Ala in 4Q9I), *Ec*MsbA (LPS in 5TV4) and BtMRP1 (Leukotricine C4 in 5UJA)), positions homologous to residues in TAP and P-gp shown to affect substrate recognition by crosslinking, allelic variants, and constructed mutations (magenta) (Srikant and Gaudet, 2019). The cyan mesh shows the cumulative density of substrates listed above. (**D**) Schematic of *Yl*Ste6 to illustrate the contiguous TM4-6 (orange, left) and TM10-12 (brick red, right) regions chosen for library construction. These TMs contain many positions highlighted in the model in c and combine to form the cytoplasm-facing cavity in the inward-open state.

To select which regions of *Yl*Ste6 to mutagenize, we reviewed mutational analysis (Geng et al., 2015) and substrate crosslinking (Loo and Clarke, 2000) on homologous type I ABC exporters like P-gp and TAP, which showed that positions that interact with transport substrates or alter substrate selectivity are present over a large part of the TMD cavity. We also aligned structures of substrate-bound exporters MRP1, P-gp, MsbA (Alam et al., 2018; Johnson and Chen, 2017; Mi et al., 2017; Szewczyk et al., 2015) to identify residues in the TMD cavity whose side chains lie within 5 Å of the transport substrate and thus could also affect substrate selectivity in homologous transporters. This combination of genetics, structural analysis, and biochemistry highlights the relevance of transmembrane (TM) helices 4-6 and 10-12, which line the TMD cavity, for substrate selectivity (Figure 2c and 2d). Substrate-interacting residues in transporter structures (TAP, P-gp, MRP1 and MsbA) form large interaction surfaces with physicochemical properties that match the cognate substrate (Srikant and Gaudet, 2019). Given the size of the pheromone substrates and a large substrate interaction surface, we expected that multiple mutations are needed to alter the substrate selectivity of Ste6. Therefore, rather than systematically exploring the effect of all single mutations in this region (Fowler and Fields, 2014; Fowler et al., 2014), we used random mutagenesis, by error-prone PCR, to build mutant libraries (each with 10^4^ to 10^5^ members) on a yeast replicating plasmid, with each clone containing multiple mutations. One set of libraries mutated TMs 4-6 (483 bp, 161 aa) and the other TMs 10-12 (486 bp, 162 aa). Together, these two regions constitute the pseudosymmetric sets of TMs that form a large part of the TMD cavity surface (Figure 2d).

We performed several rounds of selection using our autocrine system to enrich for mutants in *Yl*Ste6 that improve **a**-factor export. In the first round, we subjected the libraries to FACS and gated on autocrine signal such that only ∼1% of the control, unmutagenized *Yl*Ste6 population of cells passed the selection. Given the distribution of autocrine signal in the control populations, we expect that a single round of selection is not enough to provide sufficient enrichment of functional transporters. Our system allows for expansion of the selected cells in glucose-containing medium without autocrine stimulation followed by further rounds of FACS-based enrichment after exposure to galactose-containing medium (Figure 2b). The autocrine signal of unselected mutant libraries is generally weaker than that of cells expressing WT *Yl*Ste6 (Figure 2-figure supplement 2a, lightest blue and dashed green lines respectively). We infer that most mutations are deleterious to transport function, probably by impairing the folding or stability of Ste6, as the transporter expression signal of unselected libraries was lower than populations expressing WT *Yl*Ste6 (Figure 2-figure supplement 3b). Four rounds of enrichment led to a stronger autocrine reporter signal, corresponding to selecting a subset of the transporter library with increased **a**-factor export. There was no meaningful change in autocrine reporter signal between the fourth and fifth rounds of selection (Figure 2-figure supplement 2a). We therefore performed four rounds of enrichment with each of the mutant libraries of TM4-6 and TM10-12 and sorted 10^4^ to 10^5^ cells in each round that passed our autocrine signal gate. We plated the resulting enriched populations to yield single colonies and tested these isolated clones individually using the same flow cytometry protocol; more than 95% of these clones showed increased autocrine signaling indicating improved **a**-factor export (Figure 2-figure supplement 2b and Figure 3-figure supplement 1a). To confirm that the increased autocrine signal is due to altered transporter sequences, we isolated plasmids from the selected clones, transformed them into the ancestral version of the autocrine reporter strain. Flow cytometry on these freshly transformed revealed that the selected clones encoded versions of the transporter that produced increased autocrine stimulation. The rank order of the autocrine signal in the original clones correlates well with the signal produced by transforming the isolated plasmids into fresh recipient cells (Figure 3-figure supplement 2). In summary, the FACS-based selection allowed us to enrich a library of mutant transporters for increased activity, isolate and confirm independent clones from the enriched population and therefore connect a transport phenotype to specific clones.

### Sequencing identifies mutations across the transmembrane domain (TMD) in selected clones

We analyzed the *Yl*Ste6 mutants that had an improved autocrine signal, and presumably have increased *Sc***a**-factor transport. We isolated and sequenced a total of 245 “top” clones (of about 1500 tested clones; Figure 3-figure supplement 1) from the enriched populations from six libraries (three independent enrichments each for two regions, TM4-6 and TM10-12, see methods). We used Sanger sequencing to identify the mutations present in each clone. Sequencing these 245 clones that give high autocrine signals identified 100 unique clones (61 clones from TM4-6 and 39 clones from TM10-12), suggesting that we found most of the clones in our libraries that strongly increased autocrine signaling. Multiple mutations are present in most clones with an increased autocrine signal (Figure 3-figure supplement 3). The average number of mutations per selected clone is similar to the average number of mutations per clone in the starting libraries (Figure 2-figure supplement 3). Rather than a few hotspots composed of specific positions or small regions that might represent a specific substrate contact, we observe mutations distributed across the entire mutated region in the enriched clones (Figure 3a-b; Supplementary Figure 1 and Supplementary Figure 2). We compared the resulting distributions of the number of mutations per position to the expectation that the mutations were distributed randomly across the mutated regions in a Poisson process (Figure 3c and Table 1). There is both a higher frequency of positions with zero mutations and positions that have been mutated more often than expected from a Poisson distribution. We suspect that some conserved positions cannot be mutated without severely inhibiting transporter activity, whereas others may be especially likely to affect substrate selectivity (Figure 3a-c). We also sequenced clones from a control population enriched by gating on transporter expression and not the autocrine reporter (Figure 3-figure supplement 4 and Supplementary Figure 3). Examining transporter activity in these clones confirmed that not all mutations increase the autocrine signal (Supplementary Figure 4), demonstrating that our selection enriches for mutations that increase the specific activity of the transporter.

**Figure 3.**
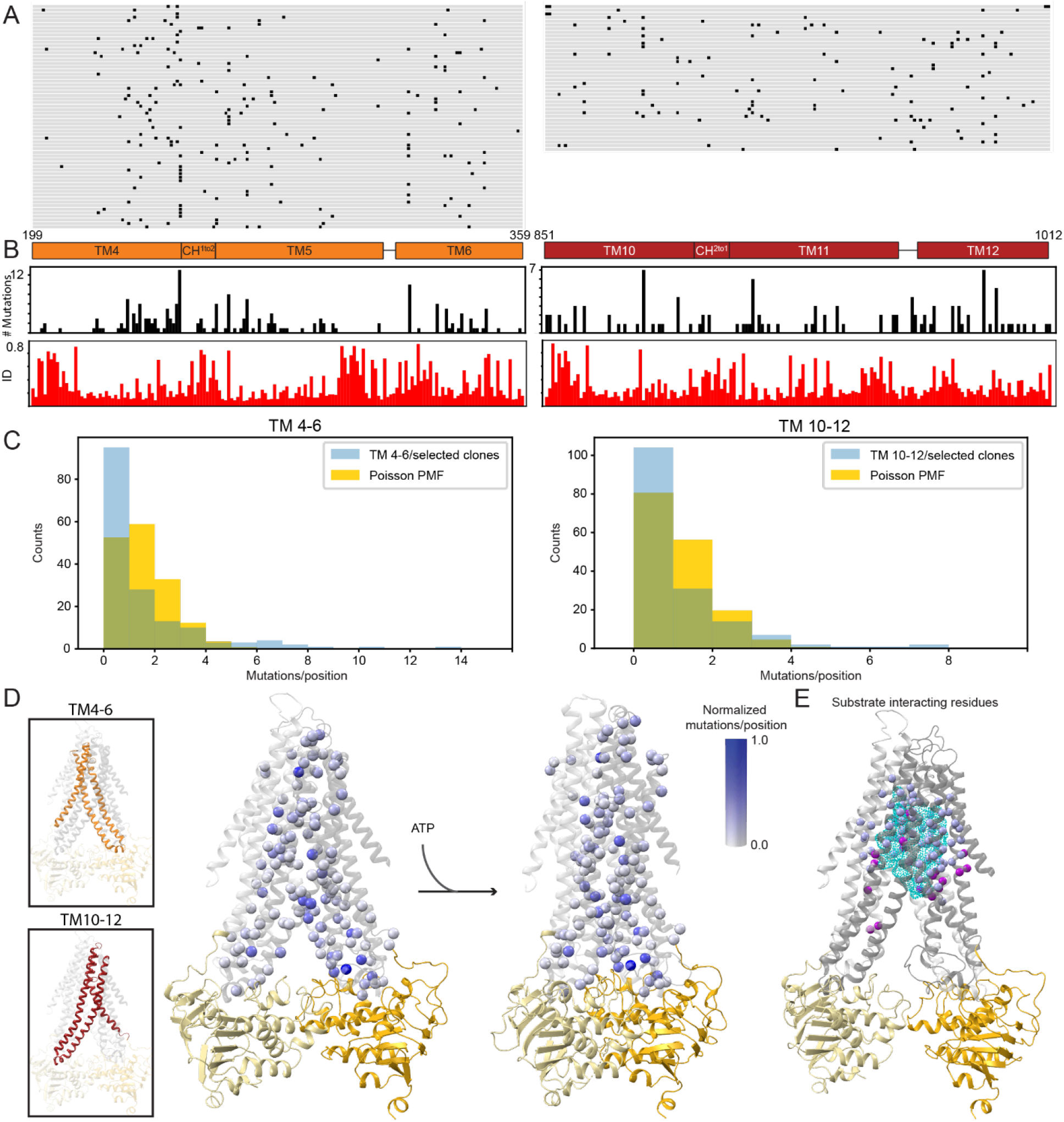
Mutations across the entire mutagenized region can increase autocrine signaling. (**A**) Enriched plasmids from the TM4-6 and TM10-12 libraries were sequenced, and the translated regions of interest are aligned to WT *Yl*STE6 with positions of non-synonymous mutations marked in black for each clone. On the left are the 61 unique clones (90 sequenced total) selected from TM4-6 libraries and on the right are the 39 unique clones (155 sequenced total) selected from TM10-12 libraries. The TM boundaries are schematized in orange (TMs 4, 5, 6 and coupling helix CH^1to2^; residues 199-359) and red (TMs 10, 11, 12 and coupling helix CH^2to1^; residues 851-1012) at the bottom of the alignment. (**B**) The number of mutations at each position summed over all unique selected clones are plotted in black bar plots for TM4-6 (left) and TM10-12 (right). The fractional sequence identity of residues in aligned fungal Ste6 sequences across the mutagenized regions TM4-6 (left) and TM10-12 (right) calculated from an alignment of 1127 fungal pheromone exporters are plotted in red bar plots. (**C**) Histogram of number of mutations per position summed over all unique selected clones from TM4-6 (left) and TM10-12 (right) libraries. The histogram of the mutations/positions (blue) has a longer tail than a Poisson probability mass function (PMF) distribution of the same number of mutations (gold). Experimental values are in Table 1. (**D**) The normalized number of mutations per position are mapped on the inward-open (left) and outward-open (right) homology models of *Yl*Ste6 with the corresponding Cα spheres colored white to blue proportional to the values plotted in Figure 4b normalized by the maximum value in each library. The insets highlight the symmetric regions TM4-6 and TM10-12 that are mutated in our libraries. (**E**) Figure 2c panel reproduced here for comparison, with light blue spheres highlighting positions proximal to substrate density, and magenta spheres highlighting positions homologous to residues in TAP and P-gp shown to affect substrate recognition by crosslinking, allelic variants, and constructed mutations.

**Table 1.** Counts of mutations per position in selected clones.

| TM4-6 clones |  | TM10-12 clones |  |
| --- | --- | --- | --- |
| No. of mutations | Positions | No. of mutations | Positions |
| 0 | 95 | 0 | 104 |
| 1 | 28 | 1 | 31 |
| 2 | 13 | 2 | 14 |
| 3 | 10 | 3 | 7 |
| 4 | 3 | 4 | 2 |
| 5 | 3 | 5 | 1 |
| 6 | 4 | 6 | 1 |
| 7 | 2 | 7 | 2 |
| 8 | 1 |  |  |
| 9 | 0 |  |  |
| 10 | 1 |  |  |
| 11 | 0 |  |  |
| 12 | 0 |  |  |
| 13 | 1 |  |  |

NOTE: This data is plotted in Figure 3c.

Plotting the number of mutations per position from the clones selected for increased autocrine signaling on homology models shows that mutations in selected clones are distributed throughout and beyond the TMD cavity surface (Figure 3d). We compared these frequently mutated positions to positions in homologous type I exporters where mutations change substrate recognition (see Figure 3e). Our selected clones identify a larger set of positions where mutations increase *Sc***a**-factor transport and could be available to evolve new transporter specificity.

### Selected clones have increased pheromone export

We performed a more detailed analysis of the effect of mutations in selected clones. We selected four clones with high autocrine signal (two each from the TM4-6 and TM10-12 libraries; Figure 4a) to better characterize the increased autocrine reporter signal. To confirm that selection for increased autocrine signal corresponds to transporters with increased specific activity, we used two additional assays: quantifying mating efficiency and the exported **a**-factor. Cells expressing each of the four selected clones show a roughly 10-fold increase in **a**-factor export relative to cells expressing unmutated *Yl*Ste6 (Figure 4b) and the expression of these mutants improves mating efficiency relative to cells expressing the WT *Yl*Ste6 (Figure 4c). These results demonstrate that the selected clones confer increased **a**-factor transport.

**Figure 4.**
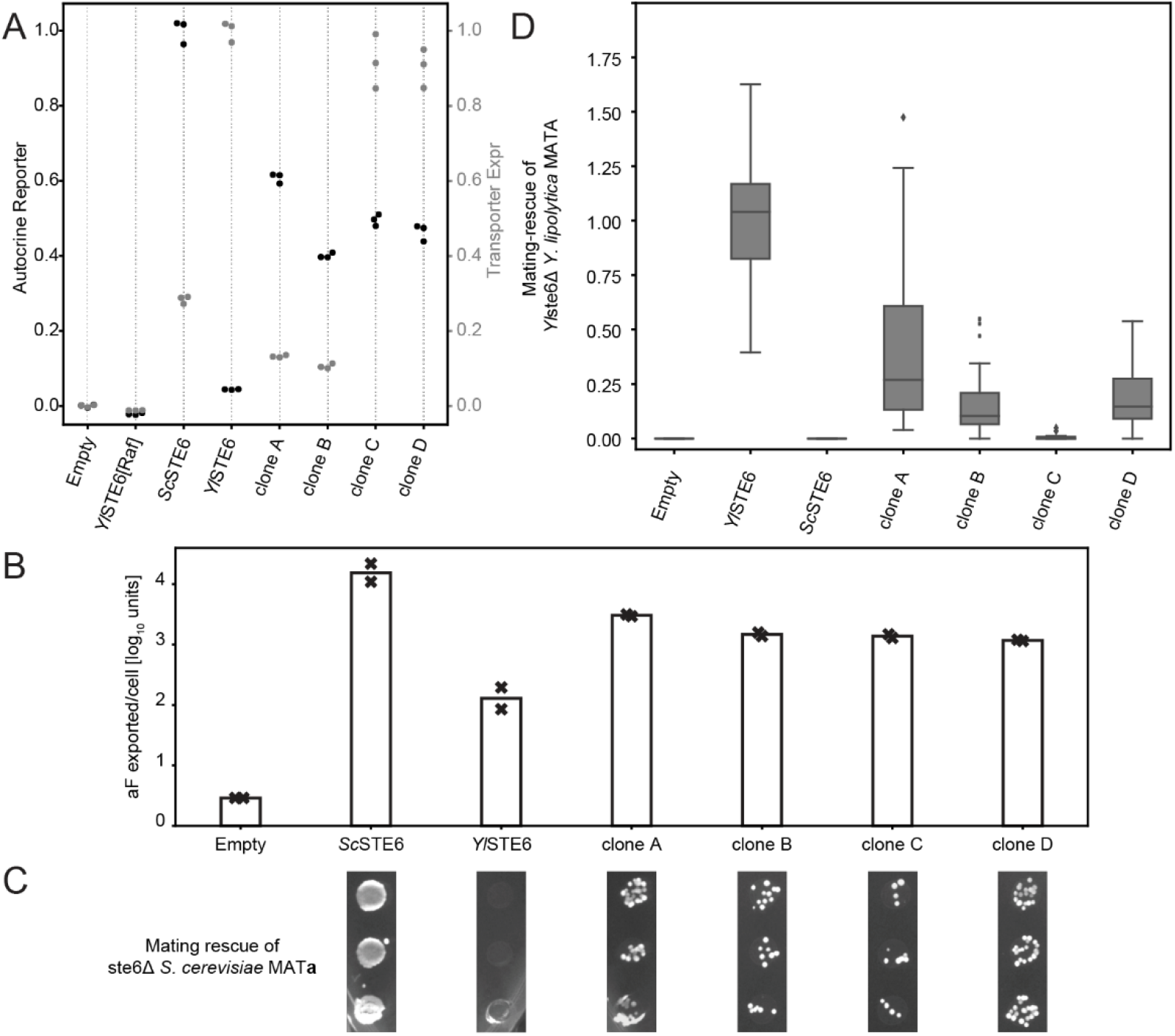
Autocrine selection produces transporters with increased *S. cerevisiae* a-factor transport. (**A**) Four clones (A and B from the TM4-6 library and C and D from the TM10-12 library) were re-introduced into naïve autocrine strains (that have not experienced FACS) to verify that increased autocrine signal is conferred by mutations in the clones. Each sample was measured by flow cytometry in biological triplicate populations (n > 25,000), and the medians of the autocrine signal (black dots) and transporter expression (grey dots) are plotted. (**B**) **a**-factor exported by populations expressing the transporter samples was measured by an end-point dilution biological assay in replicate (crosses, average denoted by bars). Selected clones A, B, C and D have increased **a**-factor export relative to WT *Yl*Ste6. (**C**) Mating rescue of *ste6Δ S. cerevisiae* strains expressing transporter samples in biological triplicate plated on diploid selective media. Clones give increased rescue of *S. cerevisiae* mating compared to WT *YlSTE6*. (**D**) Clones A, B, C and D have reduced mating efficiency in *Y. lipolytica MAT***A**-cells (equivalent to *MAT***a** mating-type of *S. cerevisiae*) when substituting for *Yl*Ste6, suggesting a reduced ability to transport *Y. lipolytica* **a**-factor. The efficiency is the number of diploids for a sample relative to the mean number of diploids formed with WT *Yl*Ste6. The box plots contain data from four different experiments, each with two biological replicates.

Mutants that increase the export of *Sc***a**-factor could act by at least three mechanisms: increasing the transport rate without altering substrate specificity, reducing transporter specificity to allow efficient transport of both *Sc***a**-factor and *Yl***a**-factor, or altering specificity to transport *Sc***a**-factor better and *Yl***a**-factor worse. To distinguish these possibilities, we assayed the *Yl***a**-factor transport function of the mutant transporters in *Y. lipolytica*. Like *S. cerevisiae ste6Δ* mutants, *Y. lipolytica ste6Δ MAT***A** (equivalent to the *MAT***a** mating type of *S. cerevisiae*) strains mate very poorly. We can restore the mating phenotype by expressing *Yl*Ste6 on a replicating plasmid, and therefore test the pheromone export activity of a mutated transporter expressed from a corresponding plasmid (Figure 4-figure supplement 1). Three of the four selected clones expressed in *Y. lipolytica ste6Δ* strongly reduce mating efficiency compared to expression of unmutated *Yl*Ste6, suggesting that these mutants have reversed the substrate specificity of *Yl*Ste6 (Figure 4d) rather than reducing substrate selectivity to allow efficient export of both pheromones.

### Individual mutations from selected clones increase pheromone export

We examined the effect of individual amino acid substitutions on *Yl*Ste6 activity and the interaction between these mutations. The four clones tested above (clone A, B from TM4-6, and clone C, D from TM10-12) each contain between 3 and 5 mutations, and we next tested the effect of each mutation in isolation. We introduced the individual mutations in *Yl*Ste6 using directed mutagenesis and measured the autocrine signal by flow cytometry with the same conditions used for selection (Figure 5a). We normalized the autocrine signal conferred by *Yl*Ste6 transporters with single mutations by baseline subtracting the WT *Yl*Ste6 signal and considered this normalized signal to be the contribution of each mutation to **a**-factor export (Figure 5-data supplement 1). Each clone has at least one mutation that increased autocrine signal when present in isolation. No singly mutated transporter is much worse than WT *Yl*Ste6, with half the mutations—C277R, Y278H, I888V, A986V, F860Y, Y940C, M1000L—being near neutral or mildly deleterious.

**Figure 5.**
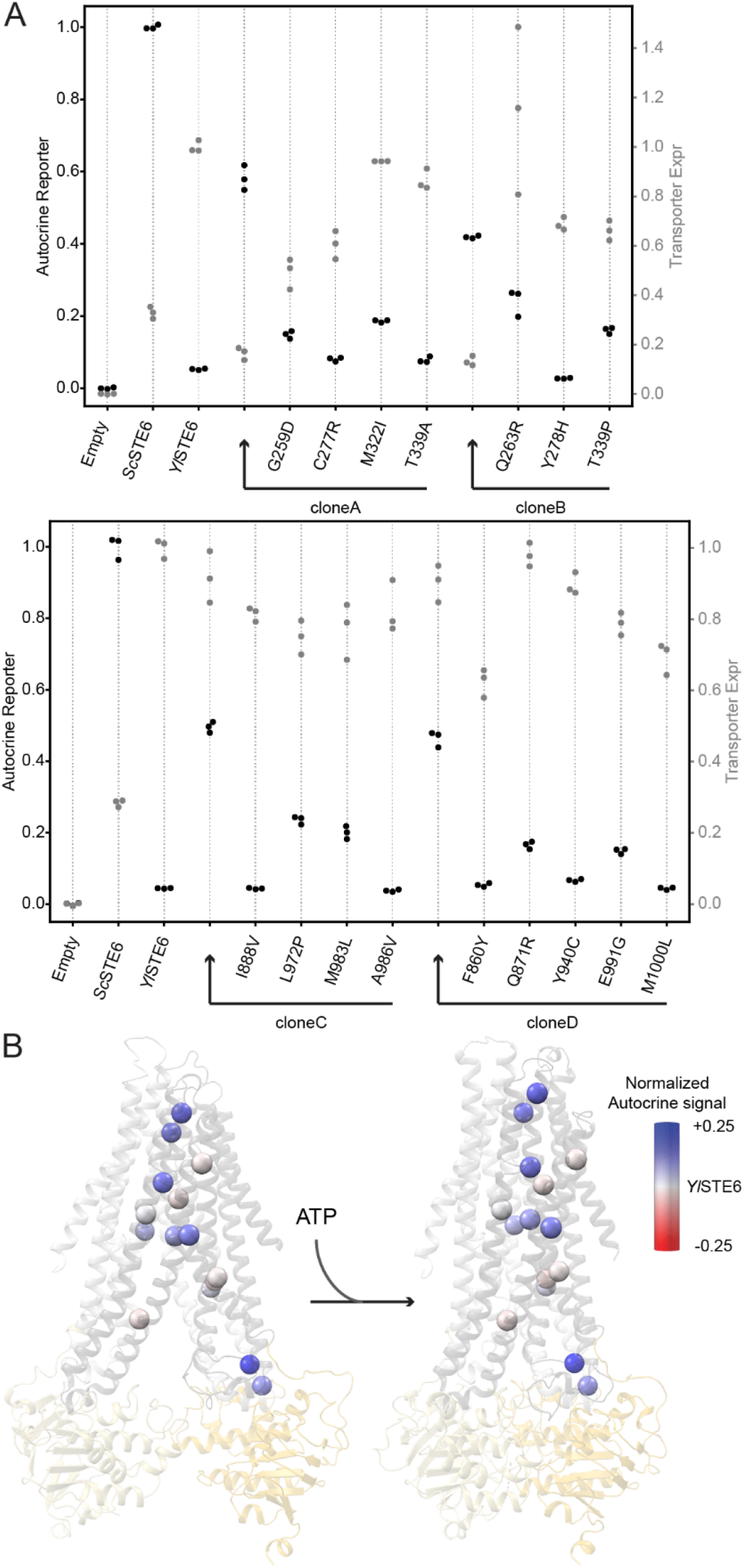
Effect of individual mutations from selected clones are neutral to positive. (**A**) Mutations in clones A, B, C and D, were tested as individual mutations by the flow cytometry-based autocrine assay, showing that many single mutations increased transport activity. Each sample was measured in biological triplicate populations (n > 25,000), and the medians of the autocrine signal (black dots) and transporter expression (grey dots) are plotted. Clones A and B from TM4-6 libraries (top) and clones C and D from TM10-12 libraries (bottom) are plotted alongside their corresponding single mutations for ease of comparison. (**B**) All mutations contained in clones A, B, C and D are plotted as Cα spheres on *Yl*Ste6 homology models and colored by normalized autocrine signal from (a). Supplementary Table 1 records the raw and normalized values used for the structural models.

Plotting the normalized autocrine signal of these single mutations on structural models, most mutations that increase the autocrine signal localize to the TMD cavity (Figure 5). This is true across all four clones, with positions of the TMD cavity buried in the lipid bilayer containing six mutations that increase the autocrine signal: M322I, T339P, Q871R, L972P, M983L, E991G. However, two strongly beneficial single mutations, G259D and Q263R, are near the coupling helices that are the structural contacts between the TMDs and NBDs. The coupling helices are important in the allosteric communication of substrate binding to the NBDs that underlies the substrate-stimulated ATPase activity that is conserved across ABC transporters (Alam et al., 2019; Pan and Aller, 2018).

### Mutations have additive effects on transport activity

We analyzed the interaction between individual mutations in a given clone to understand whether the autocrine signal of a clone depends on the specific combination of mutations. The effect of individual mutations can be positive, neutral, or negative, with the last two classes potentially hitchhiking with beneficial mutations. Doubles or triples of mutations could produce larger or smaller increases in the autocrine signal than the sum of their individual effects. In extreme cases, adding individually neutral or deleterious mutations could enhance the signal produced by other mutations. None of the mutations in the four tested clones, A, B, C and D, are strongly deleterious and therefore we are restricted to testing the effect of nearly neutral mutations in combination with other neutral or beneficial mutations.

To understand the interactions among strongly beneficial mutations, we built all possible combinations of mutations contained in clones A and C, from the TM4-6 and TM10-12 libraries, respectively, which each contained four mutations, and a subset of the combinations in clones B and D, which contained three and five mutations, respectively. We tested the combinations using the autocrine signal detected by flow cytometry (Figure 6a and Figure 6-figure supplement 1). If mutations interact additively and the autocrine signal is linearly proportional to Ste6 transport activity, the signal of multiple mutations should equal to the sum of signals of each of the mutations in isolation. Figure 6b plots the signal from all combinations of mutations we tested with the observed value on the y-axis and the sum of single mutation contributions on the x-axis. Given the error in our measurements and our uncertainty in the relationship between autocrine signaling and transporter activity, our data are consistent with beneficial mutations being additive in their contribution to the autocrine signal. We cannot exclude the possibility that the autocrine signal as a measure of transport activity may be a non-linear transform of an underlying additive property (Starr and Thornton, 2016), however, we choose to make the conservative assumption that signaling is proportional to transporter activity (Otwinowski et al., 2018). The mutations in selected clones increase the autocrine signal in an additive manner, explaining the lack of strongly deleterious mutations which would adversely affect the autocrine signal. The mating efficiency of the constructs that contain combinations of mutations from clone A or C are consistent with the measured autocrine signal (Figure 6a) and support the inference that combining beneficial mutations leads to increased **a**-factor transport. The additive nature of mutation effects highlights the fact that most single-step evolutionary paths for clones A and C are either neutral or adaptive.

**Figure 6.**
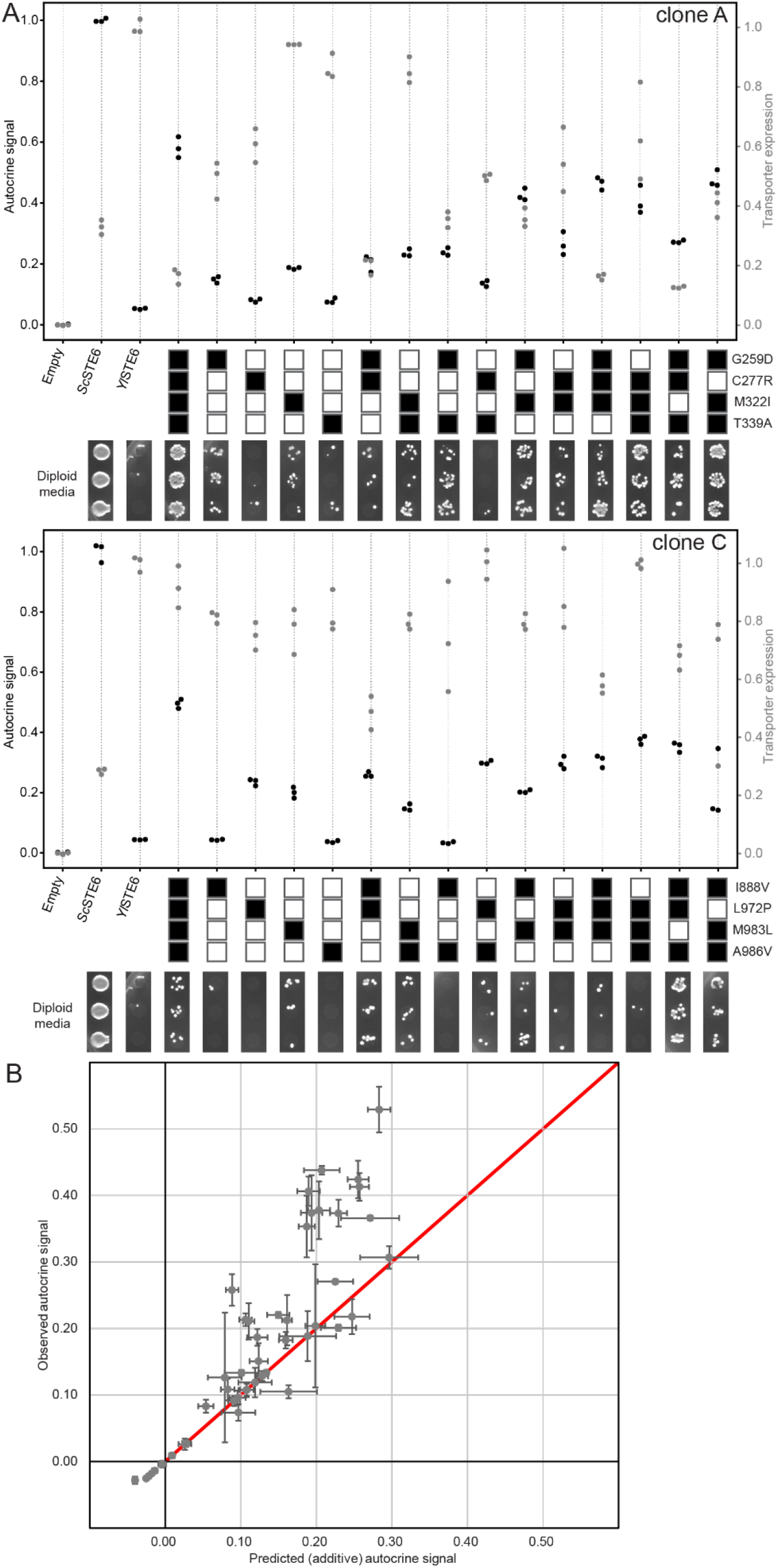
Mutations in selected clones have roughly additive effects on the autocrine signal. (**A**) Autocrine signal (black dots) and transporter expression (grey dots) of all possible combinations of the mutations found in in clones A and C were measured using the flow cytometry-based autocrine assay with biological triplicate populations (n > 25,000) for each sample. Combinations of mutations from clone A (top) and clone C (bottom) are represented by a series of boxes, with filled boxes representing the presence of a mutation. Mating activity for cells expressing *Yl*Ste6 containing each combination of mutations are displayed below the autocrine data. (**B**) Data compared to an additive model for the autocrine signal. The model, shown as a red line, is that the autocrine signal from a multiple mutant is the sum of autocrine signal of all the single mutations. The data are shown with error bars representing standard deviation of autocrine signal medians of biological triplicate populations.

When mutations are in the same clone, our enrichment for autocrine signaling selects for mutations that interact at least additively with each other. We investigated the interaction of mutations that were selected independent of each other. The clearest example is the interaction of mutations that were independently selected and lie in different parts of the protein. We therefore measured the autocrine signal of the four possible cross-library chimeras of TM4-6+TM10-12 clones, A+C, A+D, B+C, and B+D. The resulting transporters contain combinations of mutations that are beneficial but were not all selected together. As measured by flow cytometry, all the chimeras had significantly lower autocrine signal than the additive expectation (Figure 6-figure supplement 2). Thus, not all combinations of strongly contributing mutations are additive and by selecting on the combined effect of multiple mutations, our selection isolated clones that have particular combinations of mutations with high autocrine activity.

## Discussion

ABC transporters are found in all extant organisms and transport a very wide range of substrates, suggesting that their substrate specificity evolves readily. We investigated the substrate specificity of a member of the ABC transporter family that exports mating pheromones from the cytoplasm of fungal cells. Using an assay that enriches for cells with increased pheromone transport, we selected mutants of the *Y. lipolytica* Ste6 protein that can efficiently transport the *S. cerevisiae* **a**-factor. These mutants contain multiple mutations that independently and approximately additively contribute to increased pheromone transport. The multiple mutants that improve pheromone transport in *S. cerevisiae* impair transport in *Y. lipolytica* implying that they are altering rather than relaxing the transporter’s substrate specificity and are thus reversing a substrate specificity that has evolved over approximately 320 million years (Shen et al., 2018).

To understand the evolution of protein function and specificity, we must find the amino acid substitutions that alter these properties and investigate the interactions between individual mutations. But it is hard to distinguish the mutations that alter function from background variation that occurs in the course of evolution, and this challenge grows larger as the evolutionary distance between the proteins increases. Next generation sequencing, which allows high-throughput functional screening of comprehensive, single-substitution libraries of mutant proteins, elucidates the role of amino acids at individual positions in a protein (Shah and Kuriyan, 2019) and this approach can be extended to comprehensively analyze the role of combinatorial variation in a small number (≤5) of interacting positions (Podgornaia and Laub, 2015). These exhaustive screens reveal the local response of function to mutation whereas random, combinatorial mutagenesis selects for novel or altered functions that may depend on multiple mutations over a wide range of positions and thus each screen occupies a small fraction of a high dimensional sequence space. We took advantage of the well-studied molecular mechanisms of the yeast pheromone response to design a high-throughput, flow cytometry-based selection for a protein’s function by coupling its activity to the expression of a fluorescent reporter. Our approach could be easily modified to test the sequence-function relationship of other proteins in the mating pathway, including the specificity of the pheromone receptors (Di Roberto et al., 2017).

The fraction of our *Y. lipolytica* Ste6 libraries that gives increased export of *S. cerevisiae* **a**-factor allows us to make a rough estimate of how many positions in Ste6 can mutate to alter substrate specificity. In our experiments, we mutagenized a contiguous region of about 160 amino acids and obtained clones that contained between three and five mutations (Figure 3-figure supplement 3) that contributed roughly additively to improved *Sc***a**-factor export by *Yl*Ste6. To estimate the target size for mutations that produced the improved *Sc***a**-factor export we observed, we made three assumptions: that it takes mutations at three positions to improve pheromone export, that each position requires a specific mutation to improve export, and that all mutations at the third position of a codon are synonymous. In this model, there are 669,920 combinations of three out of 160 positions where mutations can occur (160-choose-3), and for each triple mutant there are six nucleotide positions (two per codon) with three alternate bases, leading to 6^3^ = 216 different combinations of non-synonymous single-base substitutions, for each group of three positions, giving a total of 1.45 x 10^8^ possible mutations. In a library of 10^5^ mutants we find about 50 independent selected clones with strongly improved autocrine signaling. This implies that there are roughly 50/10^5^ * 1.45 x 10^8^ = 72,500 three-way mutations that would satisfy our selection. If there is only one mutation at each codon that can lead to improved pheromone export, this value implies that 77 out of the 160 positions can produce such mutations (77-choose-3 = 73,150). Even if there are two possible substitutions at each position (leading to an 8-fold overcounting), the inferred number of relevant positions is only reduced to 39. Even if these calculations err by an order of magnitude, our estimate suggests that there is an enormous number of mutational trajectories that could alter the specificity of ABC transporters. The fraction of clones that show strong expression was lower in libraries produced by a higher level of mutagenesis suggesting a balance between accumulating mutations whose effects are roughly additive in providing a selective benefit without including a strongly deleterious mutation (including substitutions that prevent protein folding or function and frameshift and nonsense mutations). The cost of mutagenesis was confirmed by measuring transporter expression in unselected libraries that were mutagenized to various extents (Figure 2-figure supplement 3b). As expected, median transporter expression falls as the average mutation count of the library increases.

Type I ABC exporters are a large subfamily that share structural similarity while the specific sequence features responsible for substrate selectivity remains elusive. Type 1 exporters are involved in the drug resistance of pathogens (Fairlamb et al., 2016; Koenderink et al., 2010) and cancers (Robey et al., 2018), and mutations in several human homologs cause inherited diseases including cystic fibrosis (Borst and Elferink, 2002; Riordan et al., 1989; Schulz et al., 2017; Theodoulou and Kerr, 2015). This diversity in physiological roles highlights the variation in the substrates that different paralogs transport and suggests that understanding substrate selectivity could lead to the production of more potent and specific inhibitors of ABC transporters. Three previous types of study have given information about the positions that contribute to substrate binding of type 1 exporters: substrate-bound structures (Alam et al., 2019; Johnson and Chen, 2017; Mi et al., 2017), crosslinking of modified substrates to the exporters (Lehnert and Tampe, 2017; Loo and Clarke, 2000; Nijenhuis and Hammerling, 1996), and natural allelic variants that affect the peptides transported by TAP (Deverson et al., 1998; Geng et al., 2015; Lehnert and Tampe, 2017). Collectively, these studies identify residues located throughout the TMD cavity with a higher density near the cavity’s apex. Although informative, none of this work takes a systematic approach towards identifying residues whose mutation alters substrate specificity.

To remedy this deficit, we developed a FACS-based selection on autocrine signaling that could identify combinations of mutations that increased the transport of *Sc***a**-factor by YlSte6. We used two forms of analysis to identify positions where mutations contributed to increased transport. The first was statistical and calculated the number of positions where we saw more mutations than we would expect if the total number of mutations we observed had been randomly scattered across all positions. The second was to experimentally measure the contribution of single mutations present in selected clones, revealing that there are a number of single mutations that significantly increase the specific transport of *Sc***a**-factor. Combining these two analyses, we have evidence for 17 positions where mutations can improve export of the heterologous pheromone and 8 of these were experimentally verified (Table 2).

**Table 2.**
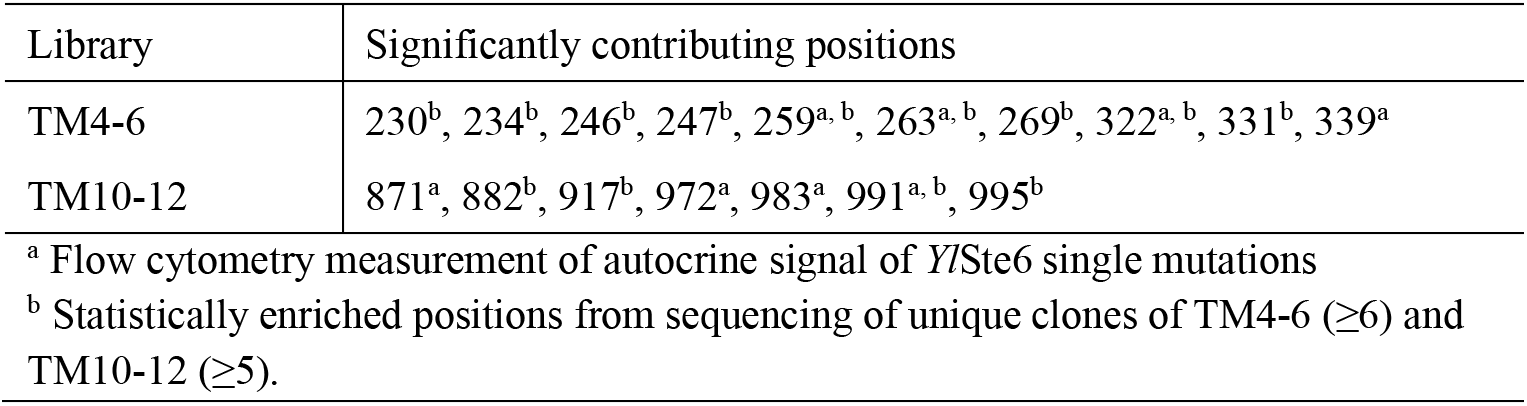
YlSte6 positions that affect Sca-factor transport.

| Library | Significantly contributing positions |
| --- | --- |
| TM4-6 | 230<sup>b</sup>, 234<sup>b</sup>, 246<sup>b</sup>, 247<sup>b</sup>, 259<sup>a, b</sup>, 263<sup>a, b</sup>, 269<sup>b</sup>, 322<sup>a, b</sup>, 331<sup>b</sup>, 339<sup>a</sup> |
| TM10-12 | 871<sup>a</sup>, 882<sup>b</sup>, 917<sup>b</sup>, 972<sup>a</sup>, 983<sup>a</sup>, 991<sup>a, b</sup>, 995<sup>b</sup> |

<sup>a</sup> Flow cytometry measurement of autocrine signal of *Y*Ste6 single mutations
<sup>b</sup> Statistically enriched positions from sequencing of unique clones of TM4-6 ( $\geq 6$ ) and TM10-12 ( $\geq 5$ ).

Among the tested mutations we see two groups of strongly contributing mutations, one at the TMD cavity buried in the membrane and the other near the coupling helices that connect to the NBD (Figure 5). Substrate-bound structures of homologous type I exporters with diverse substrates and competitive inhibitors (Alam et al., 2019; Alam et al., 2018; Johnson and Chen, 2017; Mi et al., 2017; Szewczyk et al., 2015) identify interacting residues that position substrate at the apex of the inward-open cavity (Figure 2c). Besides direct interactions that stabilize substrate binding, TM4 and TM10 undergo conformational changes in response to cognate substrate binding (Alam et al., 2019). We speculate that mutations M322I, T339P, L972P, M983L, Q871R and E991G in the *Yl*Ste6 TMD cavity contribute to substrate binding, either by direct sidechain interactions or by contributing to the flexibility of TMs needed for conformational changes. We note that four of these six mutations cause substantial changes in the sidechain. Substrate binding enhances ATP-dependent NBD dimerization, observed as substrate stimulation of ATPase activity, and is allosterically communicated by the coupling helices that form tertiary contacts between the TMD and NBD (Alam et al., 2019; Pan and Aller, 2018). Our second group of mutations, G259D and Q263R, near the coupling helix between TMD and NBD, might affect the allosteric pathway between the TMD and NBD to increase **a**-factor transport. ABC exporters contain two pairs of coupling helices that connect the two TMDs to the two NBDs. Mutations identified a role for the first coupling helix of TAP1 (between TM2 and TM3, Figure 2d) in sensing and transport of antigenic peptides (Herget et al., 2007). Although the first coupling helix identified in TAP is not contained in the mutagenized regions of our libraries, simulations of P-gp (Pan and Aller, 2018) suggest an influence of the second coupling helix (located between TM4 and TM5), which was mutated in our clones, on substrate binding and NBD dimerization. Our data suggest that mutations over a large part of the TMDs affect the transport activity of transporter in two ways: by directly affecting substrate binding and by affecting the coupling between substrate binding and ATP hydrolysis. More generally in ABC exporters, we speculate that the target size for the evolution of substrate selectivity covers a large part of the TMD and might involve both substrate binding and allostery.

We identified mutations distributed across the large surface of the TMD cavity that can potentially accommodate substrates of diverse sizes and structures. Recent cryoEM studies of different ABC exporters (Hofmann et al., 2019; Johnson and Chen, 2017, 2018) have reached consensus on the major conformational states of the substrate export cycle. The transition between inward-open and outward-open states is expected to occur via a series of microstates that lead from cognate substrate binding to conformational changes in the transporter to dimerization of the NBD, leading to opening of the substrate binding cavity to the external face of the plasma membrane. In studies on homologous transporters, mutations in the TMD cavity can either directly contact the substrate (Lehnert and Tampe, 2017), alter the flexibility of TM helices (Alam et al., 2019), affect the contacts between TM helices (Kodan et al., 2014), or potentially contribute to substrate selectivity allosterically by affecting the coupling helices and NBD interactions (Kodan et al., 2019; Pan and Aller, 2018).

We argue that the large target size and additive interaction of mutations make the evolution of substrate specificity in ABC transporters different from that of most enzymes. In contrast, work on other enzymes suggests that strong epistasis prohibits most evolutionary trajectories that would allow them to act on new substrates. As an example, analysis of β-lactamase’s ability to hydrolyze novel β-lactam antibiotics find that most trajectories are prohibited because they pass through intermediates that don’t hydrolyze the β-lactam (Weinreich et al., 2006). The effects of protein sequence context and the order of mutations further highlight the importance of both positive and negative epistasis in restricting the available trajectories for novel functions to evolve in enzymes (Miton and Tokuriki, 2016; Salverda et al., 2011). Studies of histidine kinase-response regulators and antigen-antibody protein interfaces also observe strong epistatic contributions to mutations that maintain or alter binding interfaces (Adams et al., 2019; Podgornaia and Laub, 2015). In contrast, a large number of positions can mutate to produce additive effects on the substrate specificity of ABC transporters and we propose that these relaxed molecular constraints underlie the enormous expansion of this family of primary transporters and allow them to be maintained in every branch of life.

## Supporting information

YlSte6 HomologyModels

YlSte6 ProteinAlignment

## Acknowledgements

The authors thank Michael T. Laub, Michael M. Desai, Angela Phillips, Alex N. Nguyen Ba and Thomas LaBar for critical reading of the manuscript; Claire Hartman, Jeffery Nelson and Zachary Niziolek from the Harvard Bauer Core Facility for technical assistance. We thank the members of the Murray lab and Gaudet lab for helpful discussions. SS was a Howard Hughes Medical Institute International Student Research fellow. The work was funded in part by NIH grant R01-GM120996 (to RG); and NIH grant R01-GM43987 and the NSF-Simons Center for Mathematical and Statistical Analysis of Biology at Harvard (#1764269) (to AWM).

## Author contributions

SS designed and performed experiments, analyzed and interpreted the data, and wrote the paper. RG and AWM designed experiments, analyzed and interpreted the data, and wrote the paper.

## Methods

### Strains and plasmids

All yeast strains were derived from either a *MAT***a** W303 haploid (*MAT***a**; *ade2-1*; *can1-100*; *leu2-3,112*; *his3-11,15*; *ura3-1*; *trp1-1; bud4-W303*) or a *MAT*α W303 haploid cell (*MAT*α; *BUD4*; *can1-100*; *leu2-3,112*; *his3-11,15*; *ura3Δ*) (Table 3). Strains were transformed using the LiAc-mediated chemical transformation protocol (Amberg et al., 2006). Selective markers are derived from WT versions of the *S. cerevisiae* genes, with 300-500 bp of homology flanking the marker for targeting genomic integration by homologous recombination. The *Y. lipolytica* pair of mating strains (ML16507 and ML16510) were a gracious gift from Joshua Truehart (DSM ltd) and are derivatives of the sequenced CLIB122 strain (Dujon et al., 2004). Genomic transformation was done using reported protocols (Burke et al., 2000).

**Table 3.**
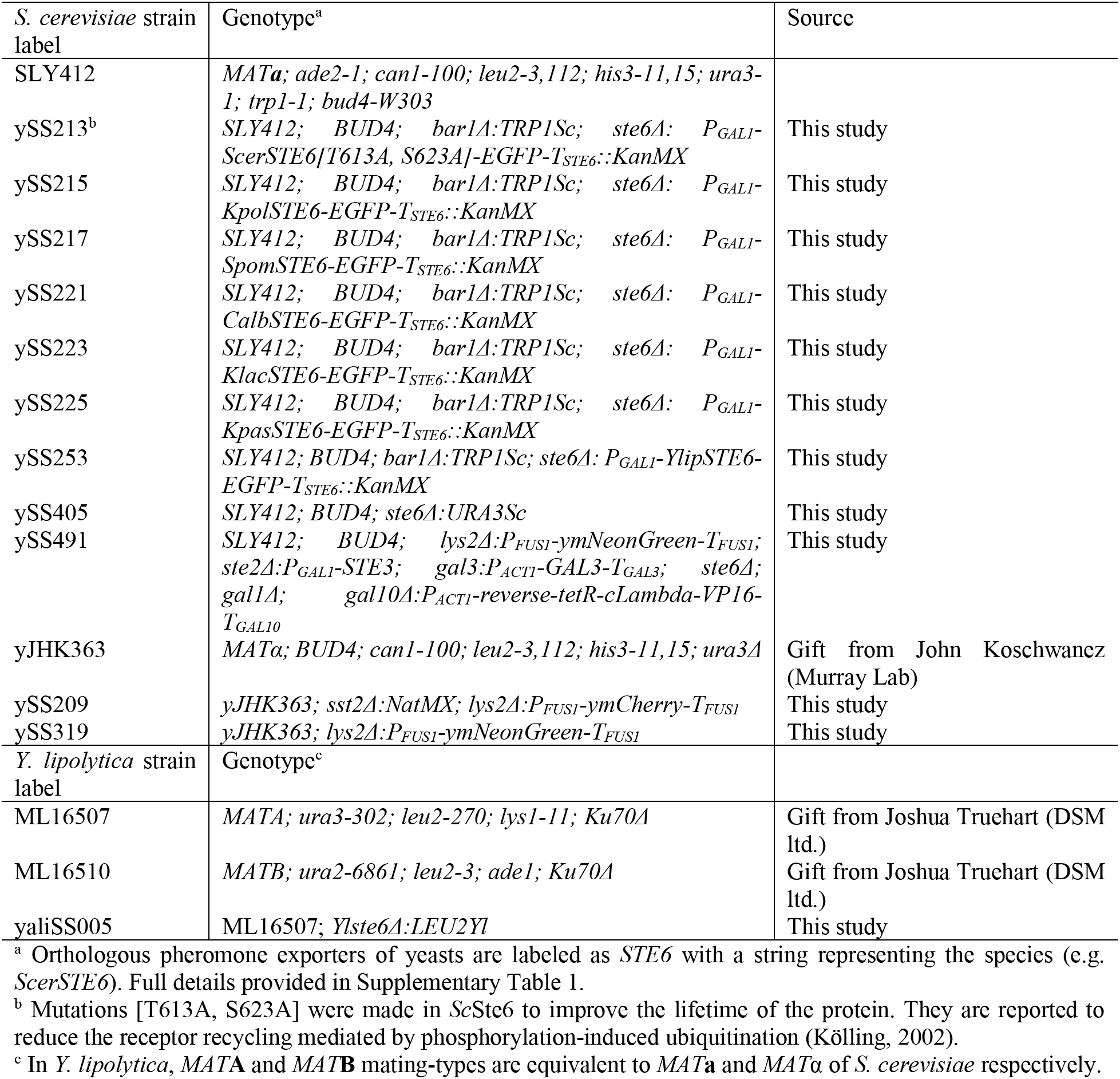
*S. cerevisiae* and *Y. lipolytica* strains used in this study.

The *S. cerevisiae* plasmid used to test the activity of heterologus pheromone transporters, pSS006, was constructed by introducing the upstream flanking region of *ScSTE6* locus (for homology), the *S. cerevisiae GAL1* promoter (*P_GAL1_*), NdeI and NotI restriction sites for ORF cloning, in-frame thrombin cleavage site, C-terminal in-frame EGFP tag (to quantify expression), *S. cerevisiae STE6* terminator (*T_STE6_*), *P_TEF1_-KanMX-T_TEF1_* (providing resistance to G418 (Thermo Fisher, Ref 11811031)), the downstream flanking region of *ScSTE6* locus (for homology), and *CEN6/ARSH6* (from the pRS41*x* plasmid series (Chee and Haase, 2012)) into pUC19 (ATCC 37254) between the BstBI and AatII restriction sites. The *S. cerevisiae* plasmid for transporter expression in the autocrine experiment, pSS021, was derived from pRS413 (*HIS3*, *CEN6/ARSH4*; ATCC 87518; (Sikorski and Hieter, 1989)), with the expression locus inserted between the XhoI and SacI restriction sites. The inserted cassette includes the *S. cerevisiae GAL1* promoter (*P_GAL1_*), NdeI and NotI restriction sites for open-reading frame (ORF) cloning, an in-frame thrombin cleavage site, a C-terminal in-frame ymKate2 tag (Shcherbo et al., 2009) to quantify expression, and the *S. cerevisiae ADH1* terminator (*T_ADH1_*). The Ste6 homologous genes were PCR amplified from genomic DNA and Sanger sequenced to confirm no differences relative to the reference available at NCBI. Ste6 homologs (Supplementary Table 1) were inserted into pSS006 and pSS021 by Gibson assembly by digesting the backbone with NdeI and NotI. *Y. lipolytica* plasmid PMB8369 was a gracious gift from Joshua Truehart (DSM ltd.), and C-terminally ymKate2-tagged transporter ORFs (as made in pSS021 above) were introduced into PMB8369 at the NotI site by Gibson assembly. All plasmid ORFs were confirmed by Sanger sequencing.

### Collection and bioassay for a-factor

**a**-factor was isolated from cultures of *S. cerevisiae* by taking advantage of its hydrophobicity as previously described (Nijbroek and Michaelis, 1998). Media used in this work are modified from Yeast extract, Peptone, Dextrose (YPD) or Complete Synthetic Medium (CSM) (Sherman et al., 1986). Briefly, overnight cultures of *MAT***a** cells (expressing different transporter homologs) grown at 30°C in YP with 3% (v/v) glycerol or CSM-His with 3% (v/v) glycerol plus 0.05% (w/v) dextrose were harvested and cells inoculated at a density of 10^8^ cells in 5 mL of collection medium (CSM with 2% (w/v) D-galactose (D-Gal), 1% (w/v) D-raffinose (D-Raf), and 0.75 µM *Sc*α-factor (peptide WHWLQLKPGQPMY, ordered from Bio-Synthesis, www.biosyn.com). Galactose induces Ste6 expression and α-factor induces maximal **a**-factor expression. The **a**-factor collection cultures were incubated in a roller drum at 30°C in a 14 mL (17×100 mm) polystyrene culture tube with inclination to maximize surface area of the tube exposed to culture. The polystyrene surface acts as an affinity resin for the hydrophobic **a**-factor secreted from cells. After 7 h, an aliquot of the culture was used to measure cell density using a Coulter Counter (for normalization of collected **a**-factor to the cell density of the cultures) and GFP fluorescence (Ex 470 nm, Em 510 nm; normalized by optical density at 600 nm, baseline subtracted with a no-transporter culture) on a SpectraMax i3. The rest of the culture was discarded, and the culture tubes were washed twice by adding 5 mL sterile water, vortexing and aspirating. The empty culture tubes were spun at 1000 g for 2 min and remaining water was aspirated. The culture tubes (caps left ON loosely) were left to dry at room temperature. One mL methanol was added to each tube and the caps were sealed tight. The tubes were briefly vortexed and left at 4°C overnight to elute the **a**-factor extract from the walls of the tube. The extract was transferred to labelled microfuge tubes and vacuum evaporated to resuspend in 40 µL methanol (a 125-fold concentration from 5 mL cultures). Two-fold serial dilutions, to a maximum dilution of 33,000-fold, were prepared from all samples in methanol and 5 µL of each dilution spotted on YPD plates (8 spots per plate, separated to avoid interference by diffusion). An overnight culture of *MAT*α-cells (ySS209) at 10^6^-10^7^ cells/mL was sprayed onto the spotted plates using an atomizer (Oenophilia, REF 900432) to generate a uniform lawn. After the plates were dried, they were incubated at 30°C overnight, and imaged. The end-point dilution of **a**-factor in the extract was the lowest extract concentration that still prevented growth of the *MAT*α-cell lawn on the spot. The **a**-factor exported per cell was determined as the fold-dilution of extract that no longer arrests cell growth divided by the number of cells in the collection culture at the end point (7 h). The data are reported in log_10_ units because the precision of measurements is restricted by serial dilution.

### *S. cerevisiae* 96-well plate mating assay

The *S. cerevisiae* mating assay was modified from a classic quantitative filter mating assay (Sprague, 1991) into a high-throughput, semi-quantitative test. *MAT***a**-cells (ySS405) containing pSS021 (*HIS3/CEN*) or pSS006 (*KanMX/CEN*) derivatives containing the transporters to be tested were pre-cultured overnight at 30°C in selective medium (2% (v/v) glycerol + 0.05% (w/v) dextrose) in a 96-deep-well block with biological triplicate colonies, while a single large culture of *MAT*α cells (ySS319) was started in YPD to use as a mating partner. The cells were pelleted, washed in water and 5 x 10^6^ cells of each mating type were mixed in a flat-bottom 96-well plate in 200 µL CSM + 2% (w/v) D-galactose and the plate was incubated at 30°C for 6 h to allow the cells to settle and form zygotes. The D-galactose is sufficient for saturating transporter induction of the *GAL1* promoter (*P_GAL1_*). Fifteen µL of 20% (w/v) dextrose stock was added to each well, the plate was sealed with breathable film and agitated for 250-300 minutes to allow zygotes to reenter the cell cycle. The samples were pelleted, medium was discarded and the cells were resuspended in 150 µL sterile water and then printed on media that selected for haploids (CSM-Lys for *MAT***a**, and CSM-Ade for *MAT*α) or diploids (CSM-Lys-Ade) and incubated at 30°C overnight. The plates were imaged, and images were processed in Fiji (ImageJ) (Rueden et al., 2017; Schindelin et al., 2012).

### Error-prone PCR mutagenesis and library construction

The error-prone PCR mutagenesis protocol reported in (Cadwell and Joyce, 1992) was modified, using various concentrations of Mn^2+^ as the mutagenic agent. Briefly, the protocol provided with Taq polymerase (NEB, Cat# M0273S) was followed while using 0.5 mM, 0.25 mM, 0.125 mM or 0.0625 mM MnCl_2_ to mutagenize TM4-6 (residues 199-359; primers oSS10_062 and oSS10_063) or TM10-12 (residues 851-1012, primers oSS10_089 and oSS10_090) (Table 4). The reactions were run over 30 cycles with 10-20 ng of template DNA (WT *Yl*Ste6 in plasmid pSS021) in a Bio-Rad S1000 thermal cycler. The number of mutations per clone was estimated by Sanger sequencing ∼10 clones from each Mn^2+^ concentration. The mutation rate increased monotonically with Mn^2+^ concentration (Figure 2-figure supplement 3). About 90 clones from a TM4-6 library mutagenized with 0.5 mM Mn^2+^ were sequenced to confirm our estimate of mutation rate. This study ran six independent enrichments of libraries, three each for TM4-6 and TM10-12. The three enrichments for each region were from independent PCR mutagenesis; two enrichments with independent libraries mutagenized with 0.5 mM Mn^2+^, the third enrichment of libraries with libraries mutagenized 0.25 mM and 0.125 mM Mn^2+^, respectively.

**Table 4.**
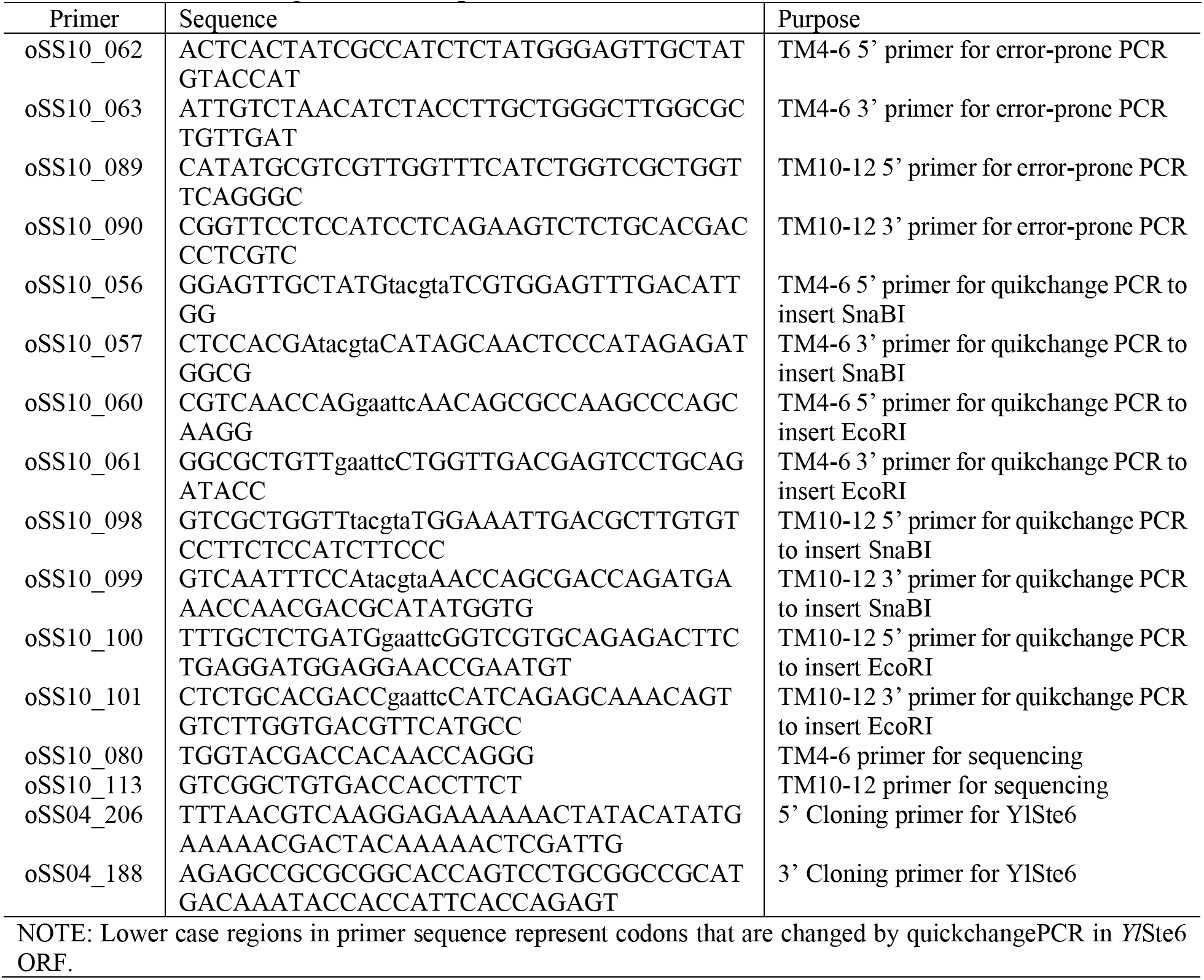
Primers used to generate error-prone PCR.

Our libraries were built *in situ* by co-transforming the mutated region and backbone linear fragments into *S. cerevisiae* and relying on homologous recombination to create circular plasmids containing mutated versions of *Yl*Ste6. Primers used for error-prone PCR are homologous to the ends of SnaBI- and EcoRI-digested backbone of *Yl*Ste6[TM4-6]-pSS021 and *Yl*Ste6[TM10-12]-pSS021. The restriction sites were introduced by a modified site-directed mutagenesis protocol (Zheng et al., 2004), where a primer pair is designed to introduce specific mutations on a plasmid by PCR. We used electroporation to transform the linear fragments into *S. cerevisiae* (Manivasakam and Schiestl, 1993), to build as large libraries as possible, with a Bio-Rad pulser set to 2.5 kV, 300 Ω, and 25 µF, which led to an effective 200 Ω sample pulsed for ∼4.1 ms and gave 10^4^-10^5^ colonies per µg of transformed DNA. The cells were plated on selective medium, washed off the plate and used as the starting population of our library. Given that our libraries cover a small fraction of the large combinatorial space of mutations in TM4-6 and TM10-12, we decided to build multiple, independent libraries for each region.

### Autocrine assay and FACS or flow cytometry

The autocrine strain (ySS491) was constructed to establish an autocrine loop to fluorescently label yeast cells that have a functional pheromone exporter. In ySS491, a *MAT***a** strain, the **a**-factor receptor, Ste3, is expressed under the control of the inducible promoter *P_GAL1_* replacing the *STE2* locus (Alvaro and Thorner, 2016; Huberman and Murray, 2013), and ymNeonGreen is expressed under the control of the pheromone-induced promoter, *P_FUS1_*, replacing the *LYS2* coding sequence (Ingolia and Murray, 2007; Poritz et al., 2001). *P_GAL1_* is converted to an inducible promoter by disabling the positive feedback loop of the galactose regulon by expressing *GAL3* from the *ACT1* promoter (Ingolia and Murray, 2007). *Sc*Ste6- or *Yl*Ste6-transformed populations were used as positive or negative controls, respectively, to measure the transporter expression (transporter-ymKate2) and the autocrine signal (ymNeonGreen) with flow cytometry. Figure 2 highlights the separation of the population clouds in a two-dimensional fluorescent color space, with the separation on the autocrine signal (x-axis) being the relevant dynamic range needed for the selections we performed. To reduce noise in batch selection and avoid paracrine signaling (exported **a**-factor influencing neighboring cells), we added 0.1% (w/v) bovine serum albumin (BSA) to the autocrine medium to effectively isolate cells from responding to **a**-factor secreted by other cells in the culture. The selection efficiency was tested by mixing cells expressing *Sc*Ste6 or *Yl*Ste6 and confirming that selecting for high pheromone-induced expression substantially enriched for cells expressing *Sc*Ste6. Due to the variation in the autocrine signal across a population of genetically identical cells, gating on high autocrine signal provided an enrichment rather than an all-or-none selection, and thus multiple rounds of sorting were advantageous.

Before selecting for autocrine signaling, the cultures were expanded in selective medium (CSM-His to select for the presence of the plasmid), with the smallest bottleneck being around 10^6^ cells, to maintain library complexity. The libraries were expanded in media with 2% (w/v) Dextrose to keep the autocrine system OFF, inoculated at 2 to 5 x 10^5^ cells/mL in 2% (v/v) glycerol plus 0.05% (w/v) dextrose to relieve catabolite repression for about 12 hrs, and then inoculated in CSM-His plus 1% (w/v) D-galactose, 1% (w/v) D-raffinose, and0.1% (w/v) BSA at 5 x 10^5^ to 10^6^ cells/mL and shaken at 30°C for 7 h. Cycloheximide (Millipore Sigma C7698) was added to 100 µg/mL to the cultures to “freeze” reporter expression of cells while sorting on an Aria flow cytometer with 488 nm and 561 nm lasers. Control populations, *Sc*Ste6 and *Yl*Ste6, were included in each experiment. Cells were sorted by setting gates for both the transporter expression and the autocrine signal. The sorting gate for transporter expression was set such that only ∼5% Autocrine OFF population (*Yl*Ste6 population in D-raffinose medium), a negative control for transporter expression passed, selecting strongly for cells with transporter expression. This was used in combination with a sorting gate for autocrine signal such that ∼1% *Yl*Ste6 population, negative control for *Sc***a**F transport passed, selecting for increased autocrine signal (Figure 2b). Library populations were sorted to collect enough events to correspond to 1-5 times the estimated library size (10^4^ to 10^5^). Collected cells were inoculated into 2% (w/v) dextrose to expand the population without selection. This enrichment was repeated over four rounds to enrich for transporters that confer higher autocrine signal.

We tested individual, selected clones, by growing them in 96-well plates and measuring their autocrine signal by flow cytometry on a High Throughput Sampler-enabled (HTS) Fortessa with 488 nm and 561 nm lasers. Biological triplicates of *Sc*Ste6 and *Yl*Ste6 controls were present in all plates for normalization.

### Clone isolation and Sanger sequencing

After enrichment, populations were plated on selective plates to isolate clones as single colonies. Clones were expanded in 400 µL CSM-His + 2% (w/v) dextrose medium in 96-deep-well blocks. The samples were pre-cultured at 1 x 10^5^ to 5 x 10^5^ cells/mL in 400 µL 2% (v/v) glycerol plus 0.05% (w/v) dextrose for 12 hr, then inoculated into a 96-well plate with 150 µL CSM-His plus 1% (w/v) D-galactose, 1% (w/v) D-raffinose, and 0.1% (w/v) BSA at 5 x 10^5^ to 10^6^ cells/mL, and shaken at 30°C for 7 h. Every plate had biological triplicates of *Sc*Ste6 and *Yl*Ste6 populations as controls to normalize the autocrine signal for each clone (described below). Plasmids were extracted from clones that showed a significant increase in the autocrine signal (relative to *Yl*Ste6 expressing cells) (Figure 3-figure supplement 1). The samples were either transformed into chemically competent *E. coli* (DH5α), for retransformation into fresh autocrine strains (ySS491), or used as template for PCR amplification of TM4-6 or TM10-12 regions for Sanger sequencing. From the isolated plasmids, 49 clones were retransformed from first two TM4-6 libraries and first TM10-12 library, and biological triplicates of these selected clones were tested in the autocrine assay. Retransforming the selected clones revealed that the autocrine signal of clones from the initial selection was a good indicator of transporter function, and physiological variation or mutations outside the regions of *Yl*Ste6 that were subjected to PCR mutagenesis were not a source of error (Figure 3-figure supplement 2).

Because the autocrine signal of isolated clones is a good predictor of transporter function, we changed our approach, for the remaining one TM4-6 library and two TM10-12 libraries, to PCR amplify the transporter ORF (oSS04_206 / oSS04_188) from the plasmid extracts from yeast clones instead of isolating plasmids by transforming these extracts into *E. coli*. The PCR reactions were Sanger sequenced with a primer for either TM4-6 (oSS10_080) or TM10-12 (oSS10_113), depending on the source library. Sequencing chromatograms were segregated based on the source library and aligned to the WT *Yl*Ste6 sequence. These alignments were processed using custom scripts (Python 3.6 with BioPython package, provided at https://github.com/sriramsrikant/FACS) to identify unique clones and calculate the number of mutations per position across unique clones. Next-generation Illumina sequencing was not used because the mutated region (roughly 480 bp) is much larger than a standard paired-end read. Illumina sequencing would thus have given us statistics on the number of distinct mutations per position but would not have provided reliable information of which mutations were linked to each other in individual clones. Given that we found 100 unique clones among the 245 sequenced by Sanger, we concluded that next-generation sequencing would not add significantly to our inferences.

### *Y. lipolytica* semiquantitative mating assay

The *Ylste6Δ MAT***A** strain (yaliSS005) was constructed from our WT *MAT***A** strain (ML16507) using a chemical transformation protocol like that used for *S. cerevisiae* (Verbeke et al., 2013). We modified a quantitative mating protocol used in (Rosas-Quijano et al., 2008) to test the mating efficiency of strains containing various transporters, carried on the *CEN*/*URA3Yl* plasmid, against our *MAT***B** partner (ML16510). Briefly, exponential cultures in YPD of the plasmid-transformed strain and its mating partner were harvested and 2.5 x 10^6^ cells of each partner were mixed in 150 µL sterile water + 0.02% (w/v) BSA. Mating mixtures were transferred onto filters (0.22 µm pore hydrophilic PVDF 25 mm membrane, Millipore Sigma Ref: GVWP02500) using a filter assembly (with the cells spreading to about 5 mm radius), and the filters (with cells) were moved onto YM mating media plates (3 g/L yeast extract, 5 g/L Bacto-peptone, 5 g/L malt extract and 20 g/L Bacto-agar) (Rosas-Quijano et al., 2008). These plates were incubated at 28°C in the dark for three days (70-74 hr). After 3 days, the filters with the mating mixtures were moved into 3 mL YP plus 2% (v/v) glycerol and 0.5% (w/v) dextrose and incubated on a roller drum at 30°C for 3 h. The cultures were transferred to microfuge tubes and sonicated to disrupt clumps, before using a Coulter counter to measure the cell density. Cells were pelleted, resuspended in water plus 0.02% (w/v) BSA and 2 x 10^7^ and 5 x 10^6^ cells were plated on diploid selective media (CSM-Lys-Ade). The mating efficiency was calculated as the number of diploid cells for the experimental samples relative to the number of diploid cells from the control mating (ML16507 + ML16510) performed on the same day (code on Github at https://github.com/sriramsrikant/). The experiment was repeated with biological replicates and plating replicates to account for the intrinsic noise of mating mixtures. Expression of transporters was estimated by flow-cytometry analysis of the ymKate2-tagged transporter, as the median of populations corrected for the background fluorescence of strains that lacked a fluorescently-tagged transporter.

### Sequence alignments and building homology models of *Yl*Ste6

The HMMER algorithm (Eddy, 2008) was used to identify homologs of Ste6 from a database of fungal proteins curated and maintained by Dr. Jim Thomas (U. of Washington). Roughly 24,000 homologous sequences were identified, down-sampled to sequences that are less than 90% identical, aligned with “hmmalign” and a sequence similarity tree constructed with “FastTree” (Price et al., 2010). The clade in the sequence tree that is monophyletic with the pheromone exporters (with known paralogous sequences as an outgroup) and with the fewest gaps accounts for 1127 sequences (Supplementary file 1). These 1127 transporters were considered orthologous pheromone exporters and aligned with “hmmalign”. The alignment was run through the ConSurf server (Ashkenazy et al., 2016; Landau et al., 2005) to identify conservation values for each position.

Homology models were built using the core transporter sequence of MRP1 (residue 314-1516) from the LTC4-bound (substrate-bound) inward-open (PDB 5UJA) and ATP-bound outward-open (PDB 6BHU) structures (Supplementary file 2,3). The structures were submitted to SwissModel (Benkert et al., 2011; Waterhouse et al., 2018) using the *Yl*Ste6 sequence as the query and the PyMOL Molecular Graphics System (version 2.2.3) (Schrodinger, 2015) was used to plot the electrostatic surface of each model. These surfaces highlight the hydrophobic band of the structure that corresponds to the transmembrane region, suggesting that the overall structure is correct, although there could be errors in the registry of helices.

