## Supplementary Material for "Selecting for altered substrate specificity reveals the evolutionary flexibility of ATP-binding cassette transporters"

Sriram Srikant, Rachelle Gaudet and Andrew W. Murray

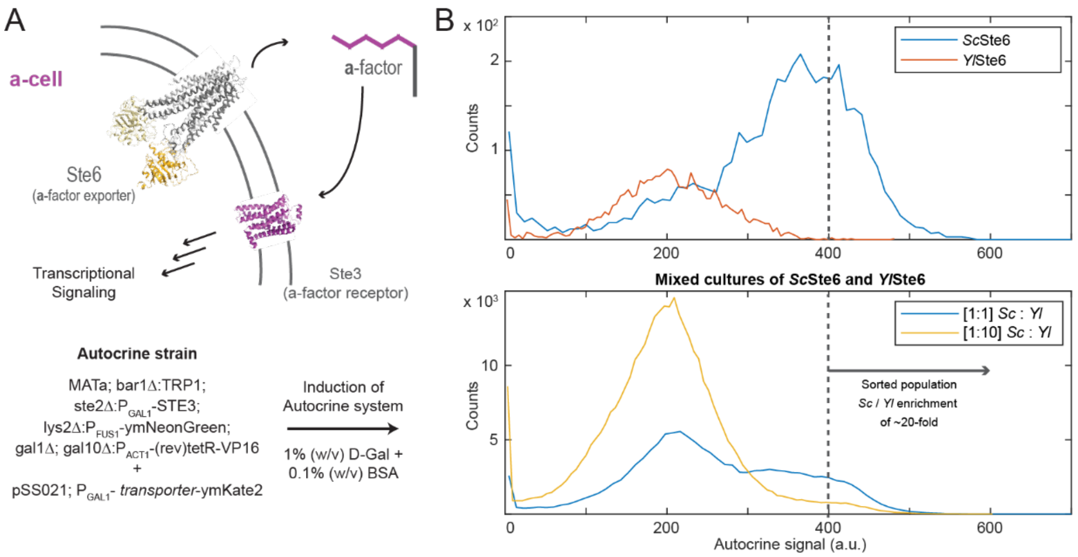


**Figure 2-figure supplement 1. Addition of bovine serum albumin (BSA) minimizes paracrine signaling and enables batch selection.** (**A**) An autocrine system was designed in **a**-cells, by expressing the **a**-factor sensitive GPCR *STE3* and a reporter for pheromone stimulation, ymNeonGreen expressed from the *FUS1* promoter (Poritz et al., 2001). The genotype of the autocrine strain is designed to induce the autocrine system with 1% (w/v) D-galactose. (**B**) Histograms of flow cytometry events in isolated *Sc*Ste6 or *Yl*Ste6 populations set a sorting gate on autocrine signal (~1% *Yl*Ste6 population) such that we can effectively enrich a 1:1 mixed population of *Sc*Ste6 *and Yl*Ste6 cells for *Sc*Ste6 by 20-fold. Thus, the autocrine system can be used for enriching transporters with increased *Sc***a**-factor transport.

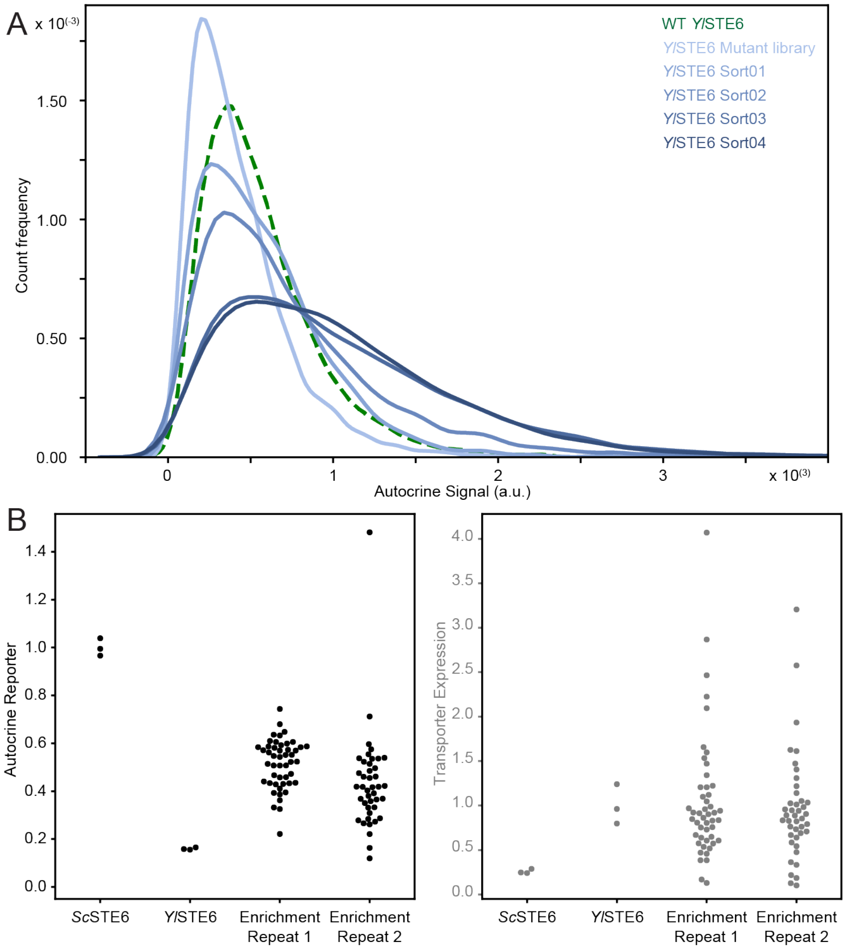


**Figure 2-figure supplement 2**. **FACS enrichment selects for clones with increased autocrine signal.** (**A**) Distributions of autocrine signal over rounds of FACS for a representative library, highlighting the selection for cells with a right shift in the distribution of the autocrine signal. The dashed green line represents a population of cells expressing WT *Yl*Ste6, while the progressively darker solid lines represent the library through rounds of FACS enrichment. (**B**) Clones isolated from enriched populations were tested individually by the flow cytometry autocrine assay to measure the clone's autocrine signal (left) and transporter expression (right). Each point in a vertical group of points represents the median measurement for a given clone.

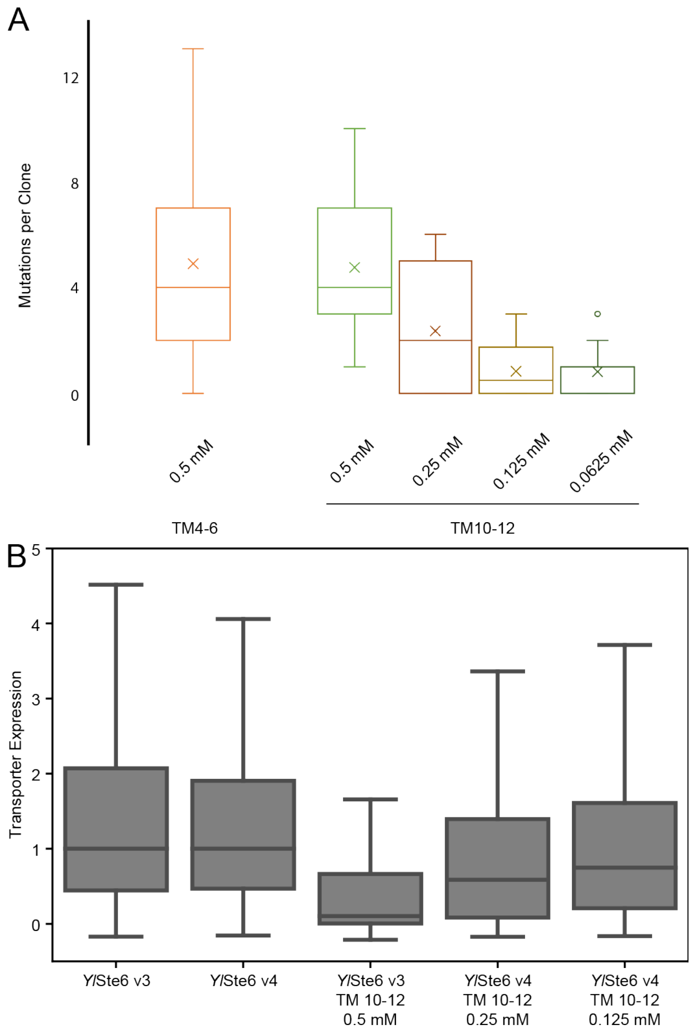


**Figure 2-figure supplement 3. Mn$^{2+}$ concentration in error-prone PCR controls the distribution of mutations per clone.** (**A**) Box-whisker plot of mutations per clone based on alignment of sequenced unselected clones with the corresponding regions of WT *Yl*Ste6, with mutations counted as nucleotide substitutions. The plot contains 83 clones for 0.5 mM Mn$^{2+}$-mutagenized TM4-6 library and 12, 11, 12 and 11 clones for 0.5 mM, 0.25 mM, 0.125 mM and 0.0625 mM Mn$^{2+}$-mutagenized TM10-12 libraries respectively. Means are plotted as crosses (X) within each distribution with outliers plotted as open circles. Box corresponds to middle quartiles and whiskers to outer quartiles. (**B**) Box-whisker plot of transporter expression levels from flow cytometry events from unselected *Yl*Ste6 mutant libraries mutagenized with 0.5, 0.25, and 0.125 mM Mn$^{2+}$. *Yl*Ste6 v3 and v4 correspond to replicate populations with WT transporter measured in independent trials with mutagenic libraries. Box corresponds to middle quartiles and whiskers to outer quartiles.

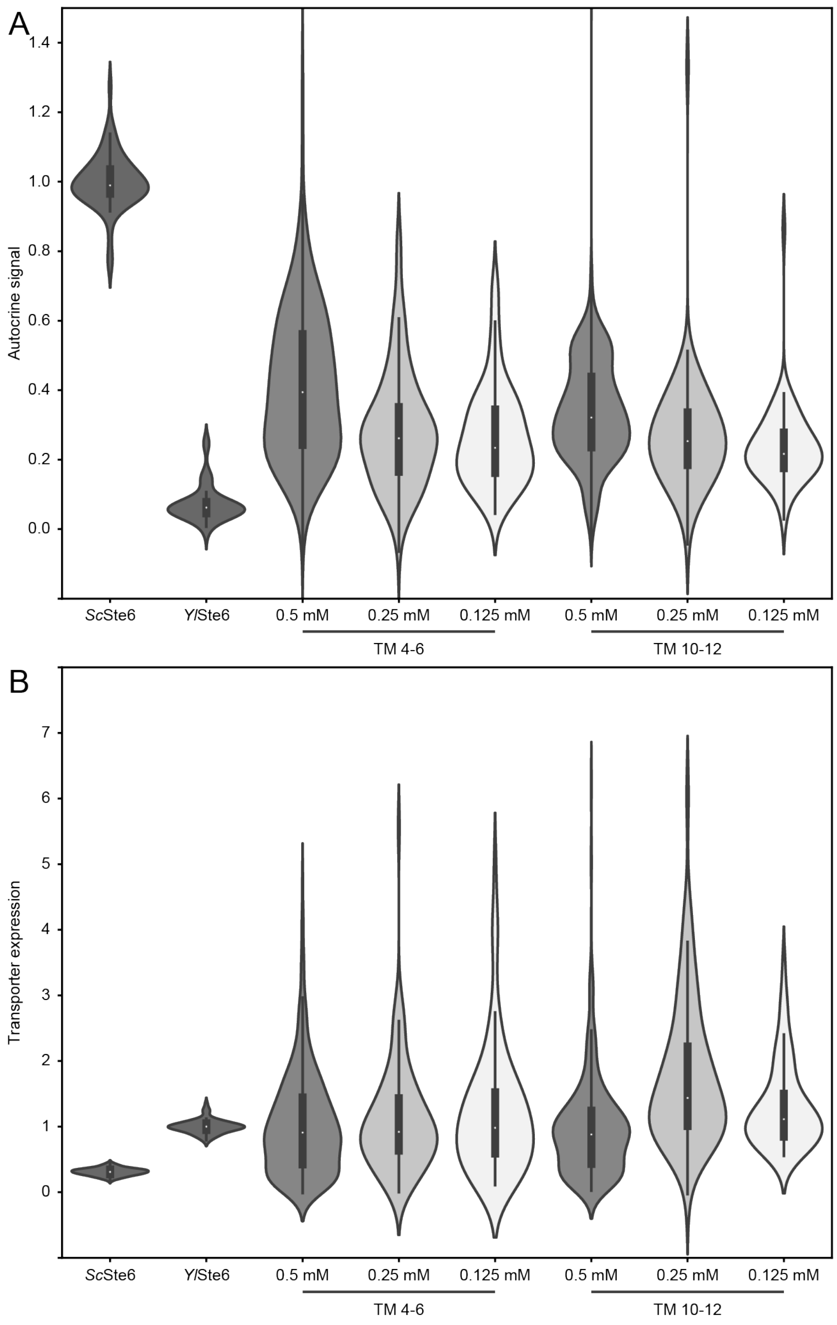


**Figure 3-figure supplement 1. Individual mutated transporter clones from enriched populations have increased autocrine signal.** (**A**) The autocrine signal and (**B**) transporter expression of isolated, selected clones pooled across replicate libraries of mutated TM4-6 (852 clones) and TM10-12 (762 clones) are consistently higher than the autocrine signal of populations expressing WT *YlSTE6* (98.0% for TM4-6 and 96.9% for TM10-12 clones are higher than WT *YlSTE6*). The signals from clones isolated from libraries mutagenized with 0.5, 0.25 and 0.125 mM $Mn^{2+}$ (shades of grey, dark to light respectively) are consistent with the fact that autocrine signal correlates with the number of (beneficial) mutations post-enrichment. The selected transporters exhibit a wide rand of expression levels. The color scheme is identical in (a) and (b).

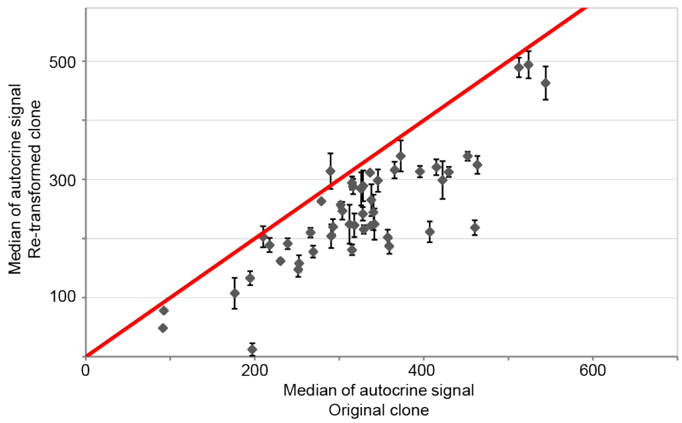


**Figure 3-figure supplement 2. Expression of mutated transporters is sufficient to confer increased autocrine signal observed in selected clones.** Plasmids were isolated from selected clones with high autocrine signal, 44 clones from TM4-6 libraries and 5 clones from TM10-12 libraries and re-transformed into the naïve autocrine strain. The autocrine signal of biological triplicates of re-transformed selected plasmids were measured by the flow cytometry-based autocrine assay. The autocrine signal of selected clones is a good predictor of the autocrine signal produced when the plasmids containing mutated *Yl*Ste6 are introduced into naïve strains that have not been subject to selection.

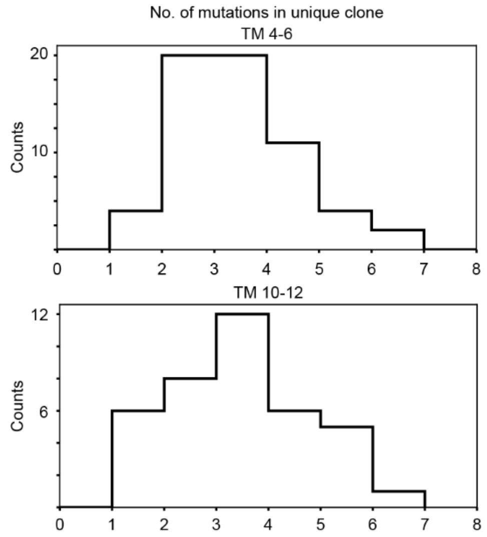


**Figure 3-figure supplement 3**. **Selected mutations contain on average 3-4 mutations** Histograms of the number of mutations per unique, selected clone from TM4-6 libraries (61 unique clones, top) and TM10-12 libraries (39 unique clones, bottom).

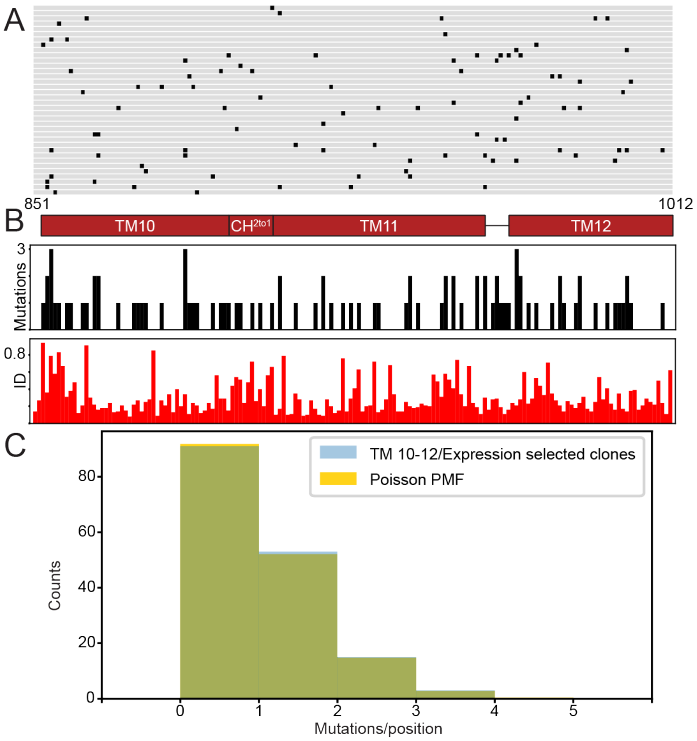


**Figure 3-figure supplement 4. Control selection for transporter expression selection yields clones with Poisson distribution of neutral mutations.** (**A**) The region of interest from 36 unique sequenced clones (89 sequenced total) selected from TM10-12 libraries based on enhanced transporter expression but not their level of autocrine signaling. The sequences are aligned to WT *Yl*Ste6 with non-synonymous mutations marked in black for each clone. The TM boundaries are highlighted in red (TMs 10, 11, 12 and coupling helix $CH^{2to1}$; residues 851-1012) below the alignment. (**B**) Like Figure 3b, the number of mutations at each position summed over all unique selected clones are plotted in black bar plots, and the fractional sequence identity of residues in aligned fungal Ste6 sequences across the mutagenized regions TM10-12 calculated from an alignment of 1127 fungal pheromone exporters are plotted in red bar plots. (**C**) Histogram of number of mutations summed over all unique selected clones. The histogram of the mutations/positions (blue) is comparable to a Poisson distribution of the same number of mutations (gold).

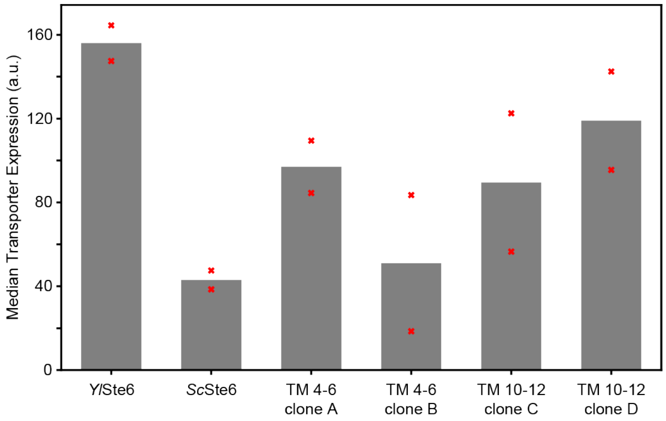


**Figure 4-figure supplement 1**. **Transporters are expressed in *Y. lipolytica* MATA cells to test for mating efficiency.** *Yl*Ste6, *Sc*Ste6, Clones A and B (TM4-6 library) and C and D (TM10-12 library) were expressed in *MAT***A** *Ylste6Δ* cells on a replicating plasmid (*CEN/YlURA3*) as used for Figure 4d. Populations were inoculated in selective media to maintain plasmids and harvested at exponential growth to measure fluorescently-tagged transport expression. Each sample was measured by flow cytometry in biological duplicate populations (n > 50,000) with medians of transporter expression plotted as red crosses, with averages of medians plotted as a bar plot.

**Figure 5-data supplement 1. Autocrine signal of singly mutated transporters**

| Sample | Clone | Autocrine signal | Transporter expression | Mutations per position | Norm. Mutations per position[a] |
|---|---|---|---|---|---|
| TM4-6 / G259D | A | 0.15 | 0.49 | 6 | 5.5 |
| TM4-6 / C277R | A | 0.08 | 0.6 | 4 | 3.6 |
| TM4-6 / M322I | A | 0.19 | 0.94 | 10 | 9.1 |
| TM4-6 / T339A | A | 0.08 | 0.86 | 4 | 3.6 |
| TM4-6 / Q263R | B | 0.24 | 1.15 | 8 | 7.3 |
| TM4-6 / Y278H | B | 0.03 | 0.69 | 2 | 1.8 |
| TM4-6 / T339P | B | 0.16 | 0.66 | 4 | 3.6 |
| TM10-12 / I888V | C | 0.04 | 0.85 | 1 | 1.4 |
| TM10-12 / L972P | C | 0.19 | 0.86 | 2 | 2.9 |
| TM10-12 / M983L | C | 0.18 | 0.89 | 3 | 4.3 |
| TM10-12 / A986V | C | 0.04 | 0.83 | 2 | 2.9 |
| TM10-12 / F860Y | D | 0.05 | 0.65 | 3 | 4.3 |
| TM10-12 / Q871R | D | 0.16 | 0.87 | 2 | 2.9 |
| TM10-12 / Y940C | D | 0.06 | 0.79 | 2 | 2.9 |
| TM10-12 / E991G | D | 0.13 | 0.65 | 7 | 10 |
| TM10-12 / M1000L | D | 0.04 | 0.67 | 1 | 1.4 |

[a] Mutations per position is normalized by the mean mutation per position of selected clones for the TM4-6 and TM10-12 libraries separately.

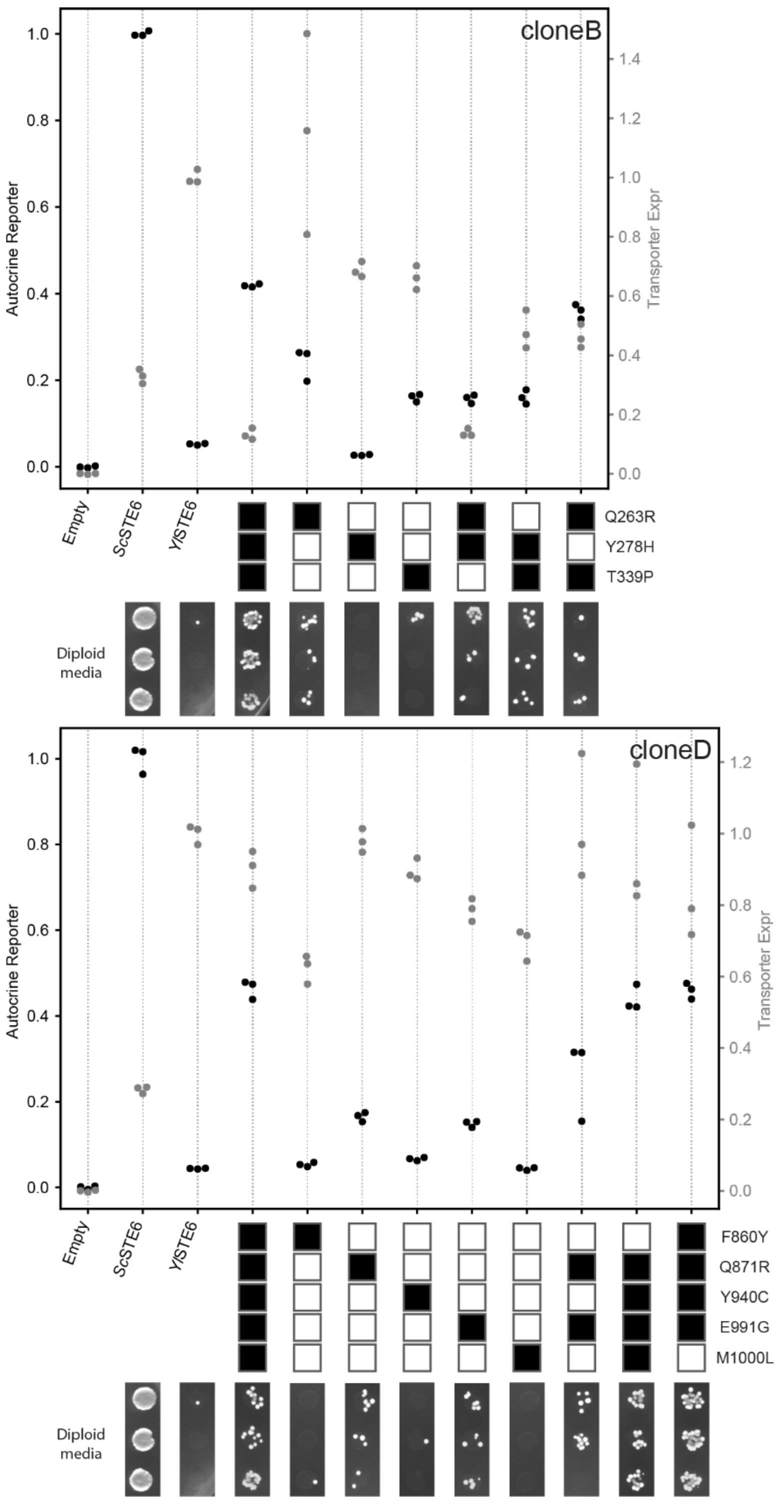


**Figure 6-figure supplement 1. Mutations from selected clones have increased autocrine signal.** Autocrine signal (black dots) and transporter expression (grey dots) of mutation combinations from clones B and D were measured using the flow cytometry autocrine assay with biological triplicate populations (n > 25,000) for each sample. Combinations of mutations from clone B (top) and clone D (bottom) are represented by a series of boxes, with filled boxes representing the presence of a mutation. Mating efficiency data for cells expressing *Yl*Ste6 containing each combination of mutations are displayed below the autocrine data. Data presented here are used to plot Figure 6b.

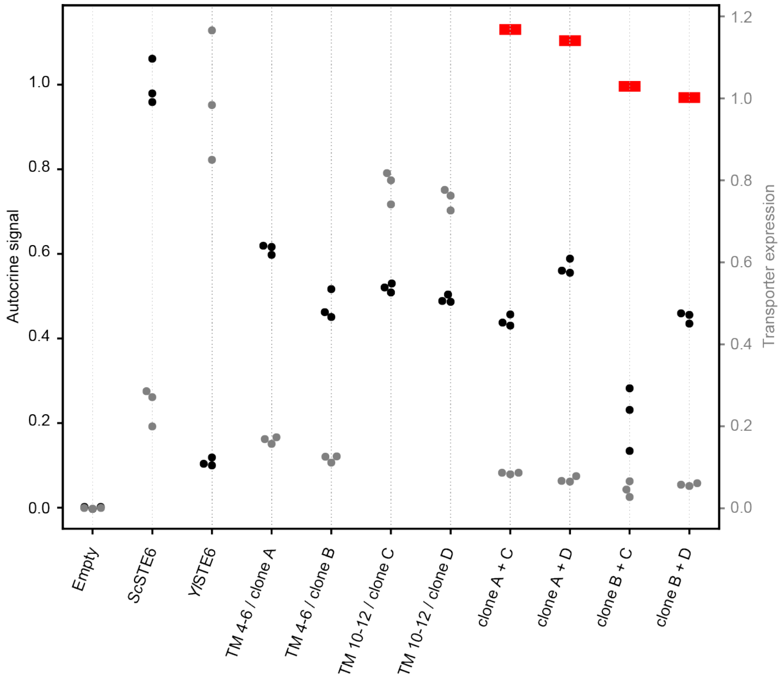


**Figure 6-figure supplement 2**. **Chimeras of selected clones do not show additive increase of autocrine signal.** Autocrine signal (black dots) and transporter expression (grey dots) of chimeras of selected clones from TM4-6 libraries (clone A, B) with clones from TM10-12 libraries (clone C, D) were measured using the flow cytometry-based autocrine assay with biological triplicate populations (n > 25,000) for each sample. For example, "clone A + C" is a chimera of the mutagenized regions of clones A (TM4-6) and C (TM10-12). The expected autocrine signal if the effects of the mutations in both regions of the protein were additive is plotted (red bars) as a reference.

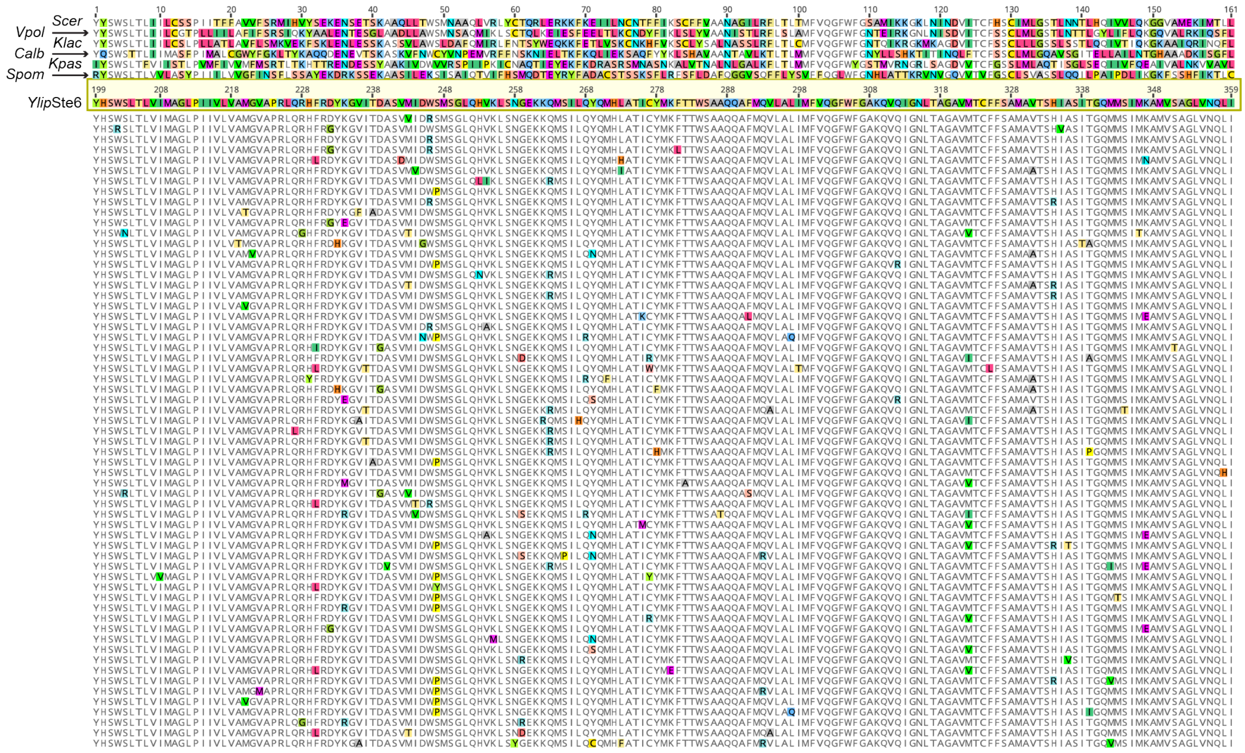

**Supplementary Figure 1. Unique clones from autocrine selection of TM4-6 *Yl*Ste6 libraries aligned to orthologous pheromone transporters.** Enriched plasmids from the autocrine selection of TM4-6 libraries were sequenced, and the translated regions of interest are aligned to orthologous pheromone transporters (Ste6, Supplementary Table 2) with positions of non-synonymous mutations with each amino-acid highlighted with a distinct color. The 61 unique clones (90 sequenced total) are ordered identical to Figure 3a.

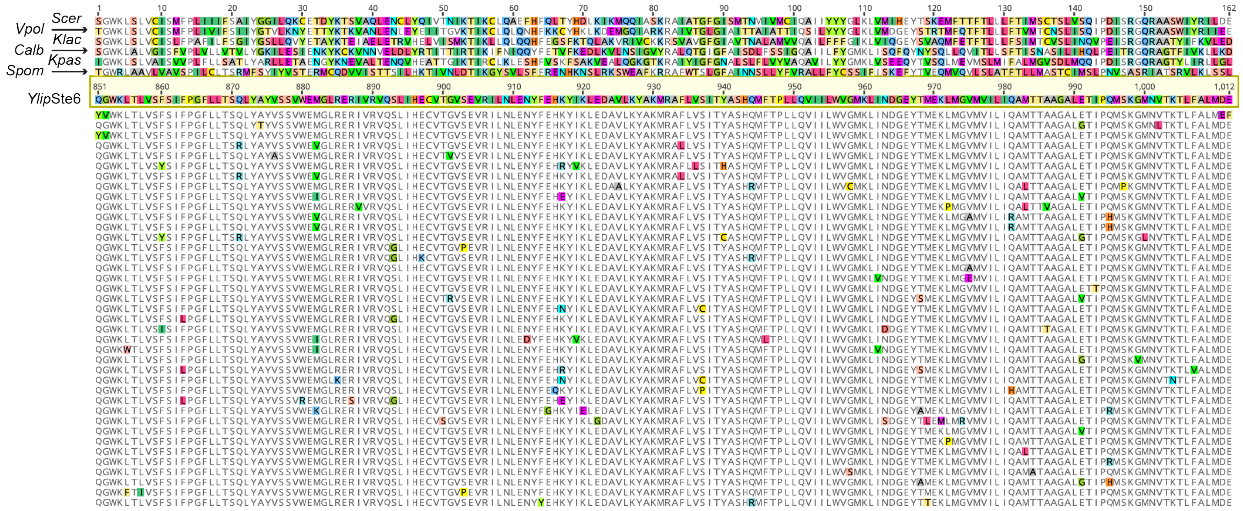

**Supplementary Figure 2. Unique clones from autocrine selection of TM10-12 *Yl*Ste6 libraries aligned to orthologous pheromone transporters.** Enriched plasmids from the autocrine selection of TM10-12 libraries were sequenced, and the translated regions of interest are aligned to orthologous pheromone transporters (Ste6, Supplementary Table 2) with positions of non-synonymous mutations with each amino-acid highlighted with a distinct color. The 39 unique clones (155 sequenced total) are ordered identical to Figure 3a.

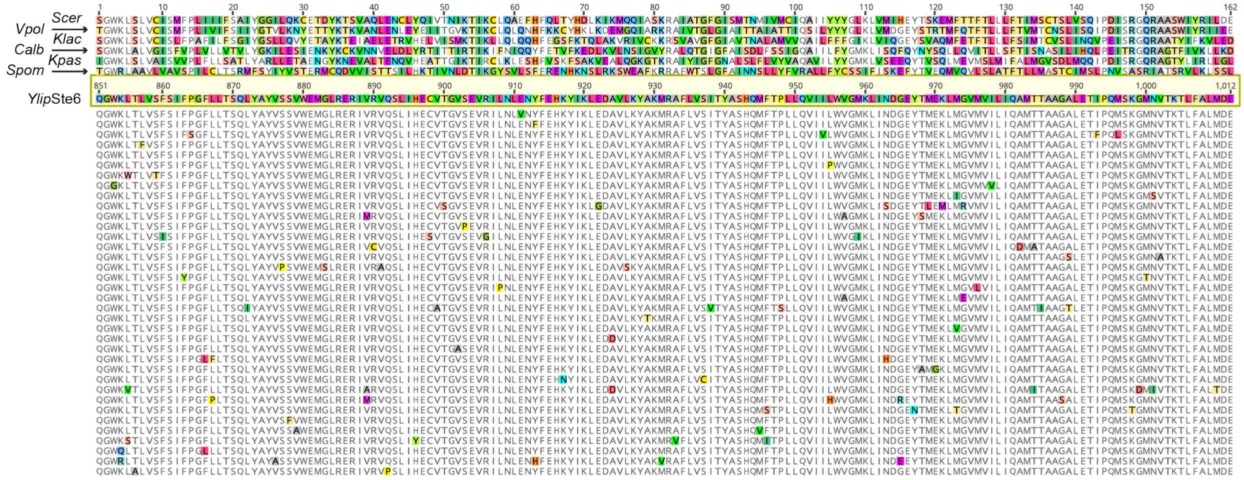

**Supplementary Figure 3. Unique clones from transporter expression selection of TM10-12 *Yl*Ste6 libraries aligned to orthologous pheromone transporters.** Enriched plasmids from the autocrine selection of TM10-12 libraries were sequenced, and the translated regions of interest are aligned to orthologous pheromone transporters (Ste6, Supplementary Table 2) with positions of non-synonymous mutations with each amino-acid highlighted with a distinct color. The 36 unique clones (89 sequenced total) are ordered identical to Figure 3-figure supplement 4a.

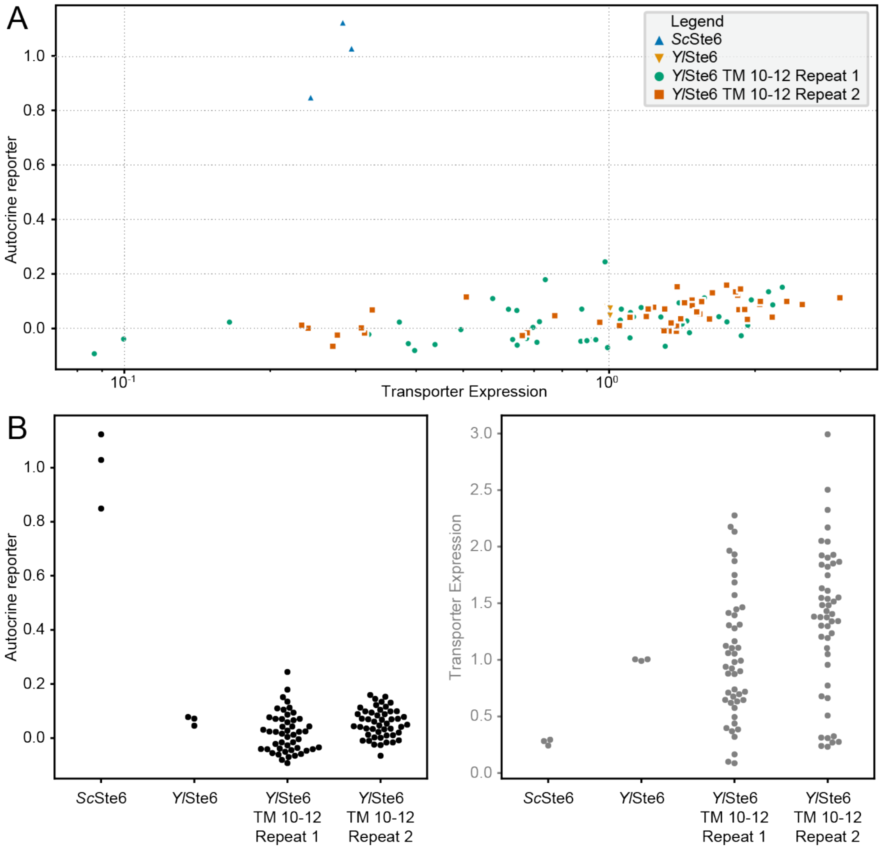


**Supplementary Figure 4. Individual mutated transporter clones from enriched populations have increased autocrine signal.** (**A**) Autocrine signal and transporter expression of isolated clones were measured using the flow cytometry autocrine assay (n > 25,000) with medians plotted for each clone. The populations were sorted by selecting on transporter expression and the resulting clones don't show a significant increase in autocrine signal. (**B**) The data plotted in (**A**) are plotted as swarm plots for autocrine signal (left; black) and transporter expression (right; grey).

| Supplementary Table 1. Homologous pheromone transporters from available species. | | |
|---|---|---|
| **Species** | **NCBI ID** | **Label** |
| *Saccharomyces cerevisiae* | *STE6* / NP_012713.1 | *ScerSTE6* |
| *Vanderwaltozyma polyspora* | Kpol1018p185 / XP_001647503.1 | *VpolSTE6* |
| *Kluyveromyces lactis* | KLLA0B14256 / XP_452165.1 | *KlacSTE6* |
| *Candida albicans* | *HST6* / KGR16750.1 | *CalbSTE6* |
| *Komagataella phaffii* | PAS_chr3_0858 / XP_002493091.1 | *KpasSTE6* |
| *Yarrowia lipolytica* | YALI0E05973 / XP_503609.1 | *YlipSTE6* |
| *Schizosaccharomyces pombe* | *MAM1* / NP_001018819.2 | *SpomSTE6* |
| NOTE: Transporter Labels are as listed in Table 3 for genotype. | | |
